# Golgi Renaissance: the pivotal role of the largest Golgi protein giantin

**DOI:** 10.1101/296517

**Authors:** Carol A. Casey, Paul Thomes, Sonia Manca, Jean-Jack M. Riethoven, Jennifer Clarke, Armen Petrosyan

## Abstract

Golgi undergoes disorganization in response to the drugs or alcohol, but it is able to restore compact structure under recovery. This self-organization mechanism remains mostly elusive, as does the role of giantin, the largest Golgi matrix dimeric protein. Here, we found that in cells treated with Brefeldin A (BFA) or ethanol (EtOH), Golgi disassembly is associated with giantin de-dimerization, which was restored to the dimer form after BFA or EtOH washout. Cells lacking giantin are disabled for the restoration of the classical ribbon Golgi, and they demonstrate altered trafficking of proteins to the cell surface. The fusion of the nascent Golgi membranes is mediated by the cross-membrane interaction of Rab6a GTPase and giantin. Giantin is involved in the formation of long intercisternal connections, which in giantin-depleted cells was replaced by the short bridges that formed via oligomerization of GRASP65. This phenomenon occurs in advanced prostate cancer cells, in which a fragmented Golgi phenotype is maintained by the dimerization of GRASP65. Thus, we provide a model of Golgi Renaissance, which is impaired in aggressive prostate cancer.

---

The Golgi apparatus is the central sorting and transportation hub involved in the posttranslational modification and sorting of cargo molecules, and it determines their subsequent delivery to appropriate compartments or to the exocytic and endocytic pathways (Nakamura, Wei et al., 2012). In mammalian cells, Golgi is a highly organized, perinuclear, ribbon-like structure, composed of stacks of flattened and elongated cisternae. In response to stress, inhibition of endoplasmic reticulum (ER) to Golgi trafficking, and treatment with or consumption of various chemicals and alcohol, Golgi undergoes remodeling, characterized by varying degrees of scattering and unstacking (Klausner, Donaldson et al., 1992, Pavelka & Ellinger, 1983, Petrosyan & Cheng, 2014, Petrosyan, Cheng et al., 2015b, Rogalski, Bergmann et al., 1984, Romero, Renau-Piqueras et al., 2015, Zaal, Smith et al., 1999). In addition, mitosis and apoptosis are also accompanied by Golgi fragmentation (Nozawa, Casiano et al., 2002, Stanley, Botas et al., 1997). Nevertheless, Golgi has a remarkable self-organizing mechanism (Misteli, 2001), and in most cases except apoptosis, Golgi returns to its classical structure and positioning as soon as cells return to a drug- or stress-free condition (Berger, Grimm et al., 1993, Lippincott-Schwartz, Yuan et al., 1989, Sweeney, Siddhanta et al., 2002, Zilberman, Alieva et al., 2011). This fact is of particular note because, despite over 100 years of Golgi research, scientists are still at a loss to explain the mechanisms of Golgi’s ability to maintain and recover harmony in such complicated structure.

Two types of proteins, residential enzymes and golgins, play a major role in Golgi function and morphology. While enzymes are responsible for the processing of proteins, golgins serve as the guardians of Golgi’s monolithic architecture and docking sites for many Golgi targeting vesicles (Appenzeller-Herzog & Hauri, 2006, Barr & Short, 2003, Pfeffer, 2001, Ward, Polishchuk et al., 2001). The function of golgins is coordinated by Golgi reassembly stacking proteins (GRASPs) (Dirac-Svejstrup, Shorter et al., 2000, Lin, Madsen et al., 2000, Lowe, Gonatas et al., 2000). We recently found that Golgi glycosyltransferases employ two different docking sites at the Golgi: giantin and the complex of GM130 and GRASP65 (Petrosyan, Ali et al., 2012a).

Giantin is the highest molecular weight (376 kDa) Golgi matrix protein. It consists of a short C-terminal domain located in the Golgi lumen (Sonnichsen, Lowe et al., 1998), where a disulfide bond connects two monomers to form an active homodimer, which is followed by a one-pass trans-membrane domain and then a large (≥350 kDa) N-terminal region projecting into the cytoplasm (Linstedt, Foguet et al., 1995, Linstedt & Hauri, 1993). This unique structure suggests that giantin is the core Golgi protein and therefore could be essential for cross-bridging cisternae during Golgi biogenesis (Linstedt & Hauri, 1993). A strong argument in defense of this hypothesis came from a study showing that during apoptosis, giantin is more stably associated with Golgi fragments than other golgins (Nozawa et al., 2002, Nozawa, Fritzler et al., 2004). In addition, the Warren group, using a microsurgery approach, developed Golgi-free cytoplasts from large African green monkey cells (Pelletier, Jokitalo et al., 2000), and these cells failed to form *de novo* Golgi. However, when cells were treated with Brefeldin A (BFA), followed by microsurgery, even a small amount of giantin in the cytoplast was sufficient to restore Golgi membranes.

Recently, we have reveled that ethanol (EtOH) treatment blocks activation of SAR1A GTPase, thus preventing COPII vesicle-mediated Golgi targeting of protein disulfide isomerase A3 (PDIA3), the chaperone that catalyzes dimerization of giantin (Petrosyan et al., 2015b, Petrosyan, Holzapfel et al., 2014). In EtOH-treated cells, Golgi is consequently disorganized and Golgi targeting of mannosyl (α-1,3-)-glycoprotein beta-1,2-N-acetylglucosaminyltransferase (MGAT1), the key enzyme of N-glycosylation, is altered (Casey, Bhat et al., 2016). In addition, we previously showed that giantin determines Golgi localization for core 2 O-glycosylation enzymes (both N-acetylglucosaminyltransferase 2/M and 1/L, C2GnT-M and C2GnT-L) (Petrosyan et al., 2012a, Petrosyan et al., 2014). Thus, different classes of Golgi residential enzymes seem giantin-sensitive: recent observation in giantin knockout (KO) models reveals one more glycosyltransferase, polypeptide N-acetylgalactosaminyltransferase 3 (GALNT3), which is down-regulated in cells lacking giantin (Nicola Stevenson, 2017). In addition, the siRNA-mediated KD of giantin results in an abnormal rate and glycosylation of some plasma membrane (PM)-associated anterograde cargo (Koreishi, Gniadek et al., 2013).

The alternative Golgi docking site is represented by GM130, a segmented coiled-coil dimer of which the C-terminal region binds to Golgi membranes preferentially through interaction with GRASP65 (Barr, Puype et al., 1997, Lesa, Seemann et al., 2000, Nakamura, Lowe et al., 1997, Petrosyan et al., 2012a). However, our previous results clearly indicate that in the absence of GRASP65, GM130 may form a complex with the dimeric form of giantin (Petrosyan et al., 2012a, Petrosyan et al., 2014). Given that in cells lacking both giantin and GRASP65, the intra-Golgi signal of GM130 is compromised (Petrosyan et al., 2012a), it is logical to assume the existence of a reserved Golgi tethering mechanism for GM130, one that in absence of GRASP65 can be realized through a direct link between giantin and GM130.

The growing evidence of literature indicates that Golgi fragmentation is the hallmark of cancer progression (Egea, Franci et al., 1993, Kellokumpu, Sormunen et al., 2002, McKinnon & Mellor, 2017, Petrosyan, 2015, Tan, Banerjee et al., 2017). It is important to note that in low-passage LNCaP cells (that represent androgen-sensitive prostate cancer [PCa]), giantin dimerization and therefore Golgi morphology is unaffected, while in androgen-restrictive PC-3 and DU145 cells, which serve as the models for aggressive PCa, giantin dimerization is impaired and Golgi is fragmented (Dozmorov, Hurst et al., 2009, Petrosyan et al., 2014). This phenotype, in turn, may play a significant role in cancer progression (Petrosyan, 2015), and therefore GRASP65, the only Golgi tethering partner for GM130 in these cells, becomes a critical protein for Golgi remodeling.

Here, we found that giantin, but not GM130 or GRASP65, is the key protein responsible for the restoration of juxtanuclear and flattened Golgi in cells recovering after treatment with BFA or EtOH. The ability to assess a Golgi 3D-structure using high-resolution microscopy enabled us to discover the sequential Golgi targeting of residential enzymes: giantin-independent proteins filled Golgi membranes during recovery, while giantin-sensitive did so only after a complete restoration of stacks. We found that giantin is involved in the formation of “long” intercisternal communications and that Golgi recovery required functions of Rab6a GTPase and non-muscle Myosin IIB (NMIIB) motor protein. In cells lacking giantin, PM targeting of proteins is altered and the length of Golgi membranes is reduced, but Golgi cisternae still form “short” connections via oligomerizations of GRASP65. Thus, we provide a model of Golgi remodeling that may shed light on the Golgi-mediated survival mechanism for cancer cells.

## Results

### Giantin dictates sequential targeting of Golgi residential enzymes in BFA washout cells

It has now been more than thirty years since BFA was employed as a model for the effective blockade of ER-to-Golgi transportation and rapid Golgi disorganization (Misumi, Misumi et al., 1986). BFA-induced Golgi disassembly is reversed upon drug washout (WO) (Berger et al., 1993, Lippincott-Schwartz et al., 1989), and BFA acts as an uncompetitive inhibitor that binds to the Arf1–GDP–GEF complex, thus blocking the activation of ARF1 and subsequent COPI coat assembly (Mossessova, Corpina et al., 2003, Peyroche, Antonny et al., 1999, Renault, Guibert et al., 2003). Indeed, loss of COPI vesicles from the Golgi is one of the earliest responses to BFA treatment (Kreis, Lowe et al., 1995). However, knockdown (KD) of COPI coatomer proteins, including β-COP, despite Golgi disorganization, does not mimic the crucial effect of BFA, i.e. Golgi membranes’ collapse and their absorption into the ER (Guo, Punj et al., 2008, Myhill, Lynes et al., 2008, Petrosyan & Cheng, 2013, Sciaky, Presley et al., 1997, Wang, Shen et al., 2010). This suggests that BFA is the drug having the potential to perform multiple tasks, especially in that it may have a direct effect on the structure of Golgi matrix proteins. The initial inference that BFA induces separation of resident Golgi proteins from matrix Golgi proteins (Hendricks, McClanahan et al., 1992, Lippincott-Schwartz et al., 1989, Seemann, Jokitalo et al., 2000) was revisited after findings indicating that glycosyltransferases and golgins were in the same tubular structures emanating from the Golgi and ER-localized BFA remnants (Mardones, Snyder et al., 2006, Puri & Linstedt, 2003, Ward et al., 2001). We previously detected that in Panc-1 cells expressing C2GnT-M tagged with c-Myc (Panc1-bC2GnT-M [c-Myc]), BFA treatment induces not only redistribution of C2GnT-M to the ER, but also reduces its level via proteasome-mediated degradation (Petrosyan, Ali et al., 2012b). This finding fits well with observations of other groups that BFA accelerates proteasome-dependent degradation of proteins (Dusseljee, Wubbolts et al., 1998, Sakata, Phillips et al., 2001). We and others have also shown that retrograde Golgi-to-ER transportation of Golgi residential enzymes is mediated by the interaction of their cytoplasmic tail (CT) with non-muscle Myosin IIA (NMIIA), and it is coordinated by the function of Rab6a (Miserey-Lenkei, Chalancon et al., 2010, Petrosyan et al., 2012b, Petrosyan, Casey et al., 2016, Petrosyan & Cheng, 2013, Petrosyan & Cheng, 2014, Sengupta, Satpute-Krishnan et al., 2015). Of interest, upon BFA treatment the content of most Golgi matrix proteins was also reduced, particularly in the case of giantin (Puri, Telfer et al., 2004). Given that giantin dimerization is required for intact Golgi morphology (Petrosyan et al., 2015b, Petrosyan et al., 2014), it is logical to assume that BFA-induced Golgi depolarization is accompanied by giantin de-dimerization. We tested this possibility in HeLa cells treated with 36 μM BFA for 1 hour, only because prolonged (up to 3 hours) treatment with BFA is known to induce apoptosis (**Movie S1**) (Guo, Tittle et al., 1998, Lee, Kim et al., 2013). As predicted, 1-hour treatment with BFA induces Golgi membrane dissolution into the ER (**Movie S2**). Furthermore, it not only reduces the level of giantin-monomer (Puri et al., 2004) but also induces its de-dimerization, which is restored following WO for another 30 min (**Fig. 1A**); detection of giantin-dimer was performed as previously reported (Petrosyan et al., 2014). Of note, the level of giantin-dimer was positively correlated with Golgi perinuclear positioning and was identical to untreated cells after 60 min of WO (data have not shown). This raises the possibility that giantin dimerization occurs during Golgi *de novo* formation.

**Figure 1.**
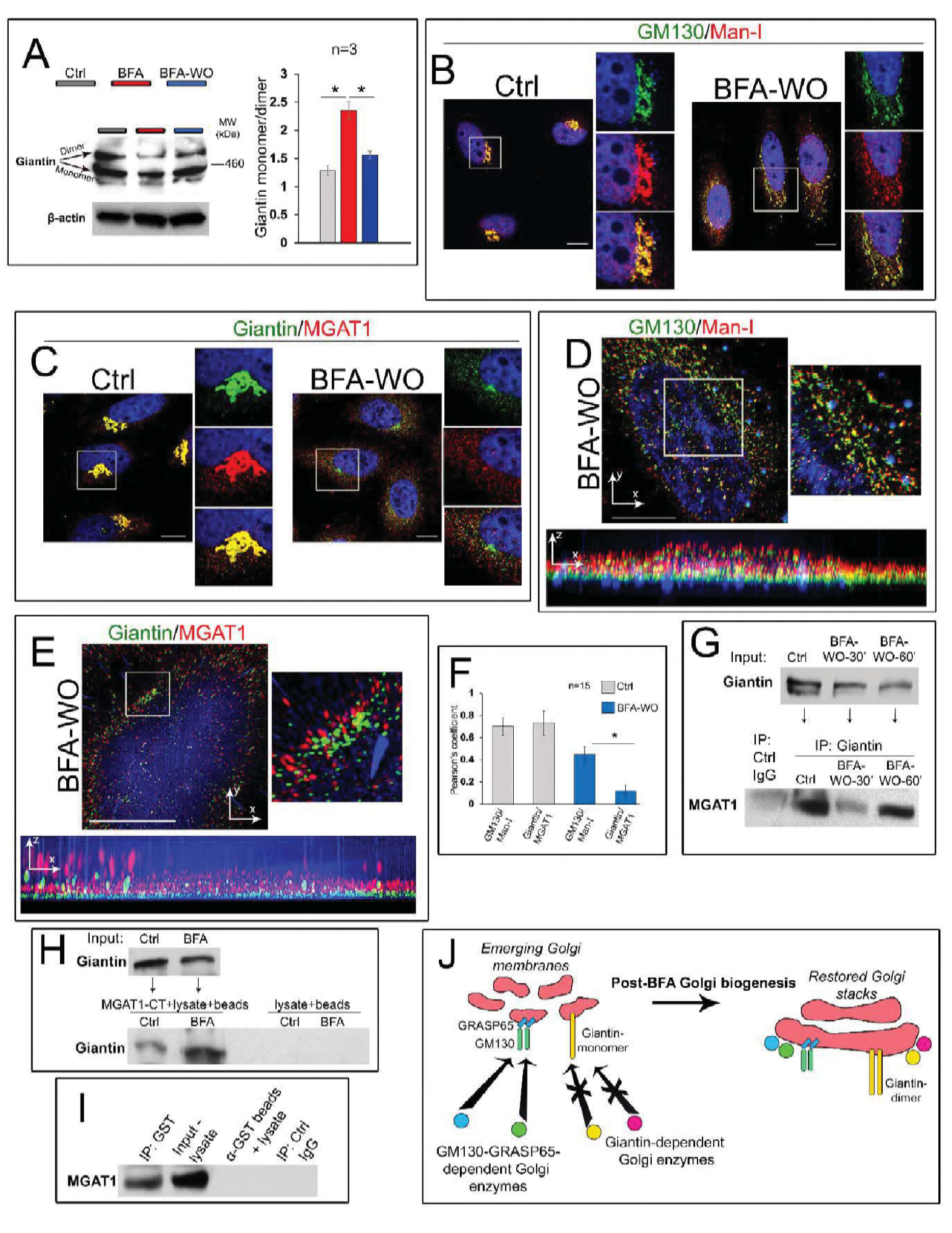
The differential Golgi targeting mechanism of enzymes during Golgi biogenesis. (**A**) Giantin W-B of the lysates of HeLa cells treated with 36 μM BFA for 1 hr or with a corresponding amount of DMSO (control), and BFA washout (WO) for another 30 min. Giantin-dimer and monomer are indicated by arrows; β-actin was a loading control. Quantification of band intensity monomer/dimer from three independent experiments. Data are presented as a mean ± SD; *, p<0.001. (**B, C**) Immunostaining of GM130 and Man-I (B), and giantin and MGAT1 (C) in control or BFA WO HeLa cells. White boxes indicate Golgi areas enlarged and shown in three channels at the right side. Nuclei were counterstained with DAPI (blue). All confocal images were acquired with the same imaging parameters; bars, 10 μm. (**D, E**) Representative 3D SIM imaging of HeLa cells after BFA WO. Cells were co-stained with GM130 and Man-I (D), and giantin and MGAT1 (E). Golgi area in the white boxes is enlarged and presented at the right side. The orthogonal views of each area are shown below; bars, 10 μm. (**F**) Quantification of Pearson’s coefficient of colocalization for indicated proteins in cells presented in D and E; n = 15 cells for each series of SD SIM imaging; results expressed as a mean ± SD; *, p<0.001. (**G**) Giantin and MGAT1 W-B of the complexes pulled down by anti-giantin Ab from the HeLa cells: control, and recovered at 30 and 60 min of BFA WO. The IP using control rabbit IgG served as a control. (**H**) Giantin W-B of the lysate (input, top panel) or the complex pulled down from the lysates of HeLa cells (treated with 36 μM BFA or a corresponding amount of DMSO) using biotinylated hMGAT1 N-terminal peptide and Dynabeads M-280 Streptavidin (low panel). Control samples are the Dynabeads incubated with cell lysate only. (**I**) MGAT1 W-B of the complexes pulled down from the lysate of non-treated HeLa cells using giantin-GST N-terminal peptide immobilized by anti-GST Ab coupled Epoxy beads. The anti-GST beads exposed to the cell lysate only and IP using control rabbit IgG are the control. (**J**) The proposed model of Golgi targeting. Enzymes that employ GM130-GRASP65 docking site are able to reach membranes during Golgi biogenesis, contrary to other Golgi resident proteins that use giantin-dimer. Targeting of the latter occurs after giantin dimerization and complete restoration of Golgi morphology.

It has been suggested that during post-BFA Golgi biogenesis, Golgi matrix proteins form a dynamic framework for subsequent delivery of glycosylation enzymes. Accumulation of these Golgi resident proteins in nascent Golgi stacks was sequential, starting from the *trans*-proteins and followed by their *cis-medial* counterparts (Jiang, Rhee et al., 2006, Kasap, Thomas et al., 2004). We recently found that the *cis*-Golgi protein, α-1,2-mannosidase (Man-I), uses GM130-GRASP65 as the Golgi docking site, whereas the next *medial-*Golgi N-glycosylation enzyme, MGAT1, uses giantin exclusively (Casey et al., 2016). Given that GM130 and giantin reside predominantly in the *cis*-and *medial*-Golgi, respectively (Petrosyan et al., 2012a, Szul & Sztul, 2011), we analyzed in BFA WO HeLa cells, colocalization of Man-I and MGAT1 with GM130 and giantin, accordingly. In untreated cells, the vast majority of Man-I and MGAT1 immunofluorescence (IF) was detected in the Golgi (**Fig. 1B and C**). In the meantime, in BFA WO cells, Man-I, but not MGAT1, was detected in recovered Golgi membranes, but the restoration of MGAT1’s intra-Golgi signal occurred only after complete recovery of the Golgi (**Fig. 1B and C, Movie S3 and S4**). To better evaluate the distribution of golgins and resident proteins in reforming Golgi membranes, we decided to employ structured illumination superresolution microscopy (SIM) to create 3D reconstructed images with a lateral resolution (~110 nm) approximately twice that of diffraction-limited instruments. The calculated Pearson coefficient of colocalization confirmed that, contrary to Man-I, MGAT1 was segregated from the emerging Golgi membranes (**Fig. 1D-F**). This is echoing the results we obtained recently in hepatocytes: giantin siRNA-mediated depletion reduced the Golgi signal for MGAT1, but not for Man-I (Casey et al., 2016). Next, we detected that giantin Ab was able to pull-down MGAT1 from the lysate of control HeLa cells. In the meantime, only marginal fraction of MGAT1 was pulled down in cells recovered for 30 min after BFA WO; however, after 60 min of recovery, when compact Golgi appears in most of the cells, the fraction of giantin-associated MGAT1 was significantly higher (**Fig. 1G**).

This finding raises the possibility that MGAT1 directly interacts with the cytoplasmic domain of giantin. Previously, to evaluate the interaction of Golgi residential proteins with NMIIA, we employed the N-terminal biotinylated peptides representing CT of different glycosyltransferases(Petrosyan et al., 2012b, Petrosyan & Cheng, 2013). Here, we used the same tool to examine whether endogenous giantin interacts with a biotin-MLKKQS peptide that matches CT of MGAT1. As shown in **Fig. 1H**, this peptide was able to pull-down giantin from the lysate of HeLa cells, both control and BFA-treated, indicating *inter alia* that despite the fact that BFA induces giantin dedimerization, it does not induce significant conformational changes in giantin-monomer, at least in terms of its potential binding to MGAT1.

Next, we hypothesize that the MGAT1 binding domain of giantin lies within its N-terminal non-coiled-coil area (UniProtKB - Q14789). To examine this plausibility, we employed the recombinant human N-terminal GST-giantin fusion peptide (3-92 aa). This peptide is successfully immobilized by anti-GST Ab coupled Epoxy beads (Dynabeads M-450), according to the GST Western blot (W-B) (data not shown). Then, the beads containing giantin-GST was incubated (overnight, at 4ºC) with the cell lysate of non-treated HeLa cells. The anti-GST Ab coupled Epoxy beads incubated with lysate only, as wells as the IP by non-specific IgG were served as a control. W-B analysis of the lysate and pull-down fractions revealed that substantial amounts of endogenous MGAT1 were pulled down by giantin-containing beads, but not by beads alone (**Fig. 1I**). Altogether, data from Figs. 1G-I indicate that CT of MGAT1 is able to interact directly with the N-terminal of giantin.

In PCa cells, we observed another example of differential Golgi targeting of O-glycosylation enzymes. As we described previously, Golgi transportation of C2GnT-L requires giantin and intact Golgi morphology, whereas Golgi-directed trafficking of the Core 1 synthase (C1GalT1, *cis-*Golgi) and β-galactoside α-2,3-sialyltransferase-1 (ST3Gal1, *medial-trans-*Golgi) is GM130-GRASP65-dependent and does not require a compact Golgi structure (Petrosyan et al., 2014). In non-treated, low passage LNCaP (c-28) cells, all three enzymes were localized to the Golgi, and predictably, BFA treatment induced their relocation to the ER (**Fig. S1A**). In BFA WO cells, both C1GalT1 and ST3Gal1 were returned to the membranes as soon as Golgi reformed back. However, the pool of C2GnT-L on the Golgi was very faint (**Fig. S1A and B**), echoing the result of MGAT1 in HeLa cells. Thus, it seems that the delayed Golgi targeting of MGAT1 and C2GnT-L is associated with incomplete giantin dimerization and Golgi reconstruction. These data suggest that the genesis of Golgi architecture and enrichment of resident proteins do not necessarily coexist simultaneously. Rather, we propose a simple model illustrating that giantin-sensitive N- and O-glycosylation enzymes are packaged in their appropriate site only within a compact and perinuclear Golgi (**Fig. 1J**).

### Giantin, but not GRASP65 or GM130, is a driver of Golgi biogenesis

These findings led us to the question whether giantin is the critical protein in the formation of juxtanuclear Golgi. To answer to this, we performed in HeLa cells the individual siRNAs (containing the pools of three to five target-specific siRNAs) depletion of giantin, GM130, and GRASP65, followed by treatment with BFA and its WO. We found that KD of GM130 or GRASP65 does not alter Golgi reassembly; however, in cells lacking giantin, Golgi remains disorganized, even though multiple Golgi fragments intend to concentrate around the nucleus (**Fig. 2A, B, and D**). Quantitatively, depletion of giantin results in almost complete loss of ability to restore compact and perinuclear Golgi, but in GM130- or GRASP65-depleted cells, the number of Golgi with this phenotype was only marginal and identical to control (**Fig. 2C**). To validate the universality of this screen, we repeated the same experiment in Panc-1 (**Fig. S2A**), A549 (**Fig. S2B**), and LNCaP (c-28) (**Fig. S2C**) cells and found that restoration of compact Golgi was significantly impaired in all types of cells lacking giantin.

**Figure 2.**
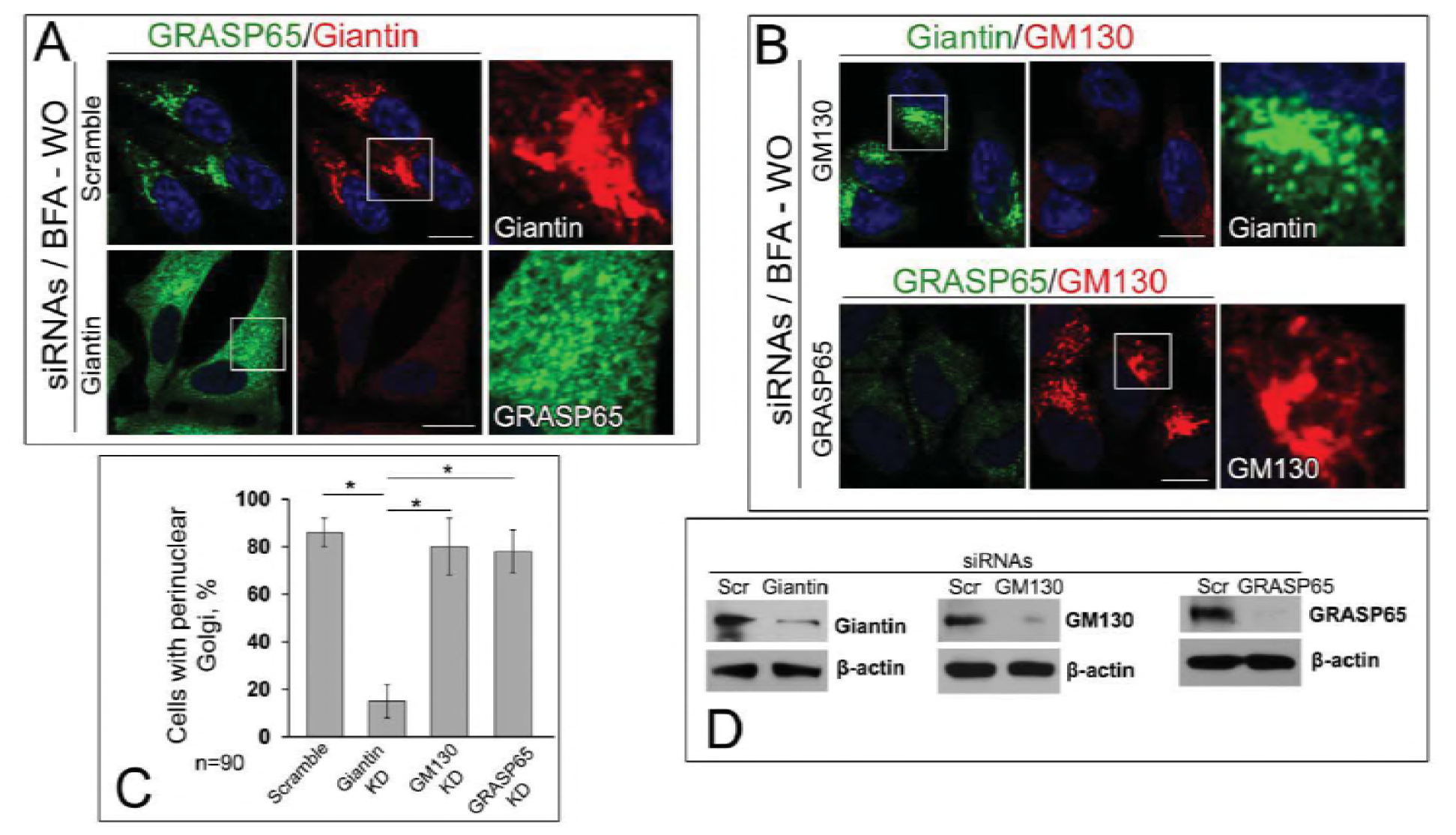
Giantin is necessary for the restoration of compact Golgi upon BFA washout. (**A-B**) Confocal immunofluorescence images of Golgi were collected in HeLa cells pretreated with different siRNAs for 72 h followed by exposure to 36 μM BFA for 60 min and then WO for 30 min. The combination of GRASP65+giantin immunostaining was used for cells treated with scramble or giantin siRNAs; giantin+GM130 and GRASP65+GM130 for cells treated with GM130 and GRAP65 siRNAs, respectively. Images in the white boxes are enlarged and displayed as either green or red channel at the right side. Nuclei counterstained with DAPI (blue); bars, 10 μm. (**C**) Quantification of cells with perinuclear Golgi as shown in A and B; n = 90 cells from three independent experiments, results are expressed as a mean ± SD; *, p<0.001. (**D**) Giantin, GM130, and GRASP65 W-B of lysates of HeLa cells treated with corresponding siRNAs; β-actin was a loading control.

### The mutuality of Rab6a and giantin

We have recently shown that giantin and NMIIA compete for Rab6a: in cells treated with EtOH, giantin de-dimerization was accompanied by the loss of its link to Rab6a. In the meantime, we observed the enhanced complex between Rab6a and NMIIA, which creates a force for Golgi disassembly (Petrosyan et al., 2016). We also reported that in PC-3 and DU145 cells, Golgi morphology appears disorganized, however, treatment with NMIIA inhibitor Blebbistatin or transfection with NMIIA siRNAs converts fragmented Golgi to the compact structure (Petrosyan et al., 2014). This Golgi “metamorphosis” required the complex between Rab6a and giantin. To test whether this phenomenon is applicable to the post-BFA Golgi biogenesis, we performed a series of experiments. First, in LNCaP cells recovered after BFA, we detected less NMIIA but more giantin associated with Rab6a than in cells after 60 min treatment with BFA (**Fig. 3A**). Second, we confirmed this observation by detecting colocalization of Rab6a with giantin but not NMIIA in the partially reconstructed Golgi of LNCaP cells after BFA WO (**Fig. 3B and C**). Third, when we performed life cell imaging of BFA WO HeLa cells that co-expressed giantin-GFP and Rab6a-RFP, the multiple giantin-Rab6a colocalizing punctae was seen in restored perinuclear structures (**Movie S5**).

**Figure 3.**
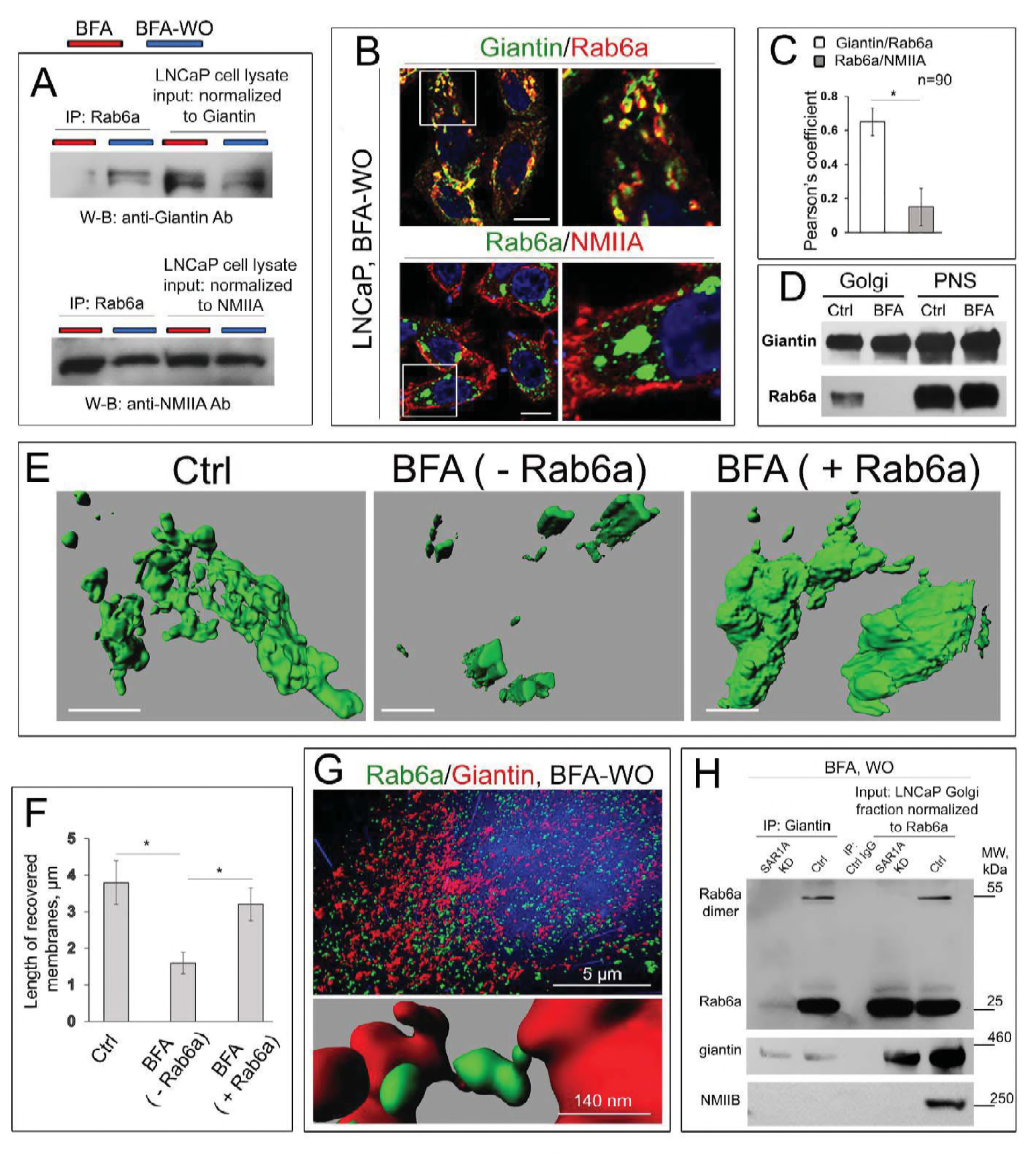
Rab6a-mediated Golgi biogenesis in LNCaP cells after BFA WO. (**A**) Giantin (top panel) and NMIIA (low panel) W-B of complexes pulled down with anti-Rab6a Ab from lysates of LNCaP cells treated with BFA for 60 min and then 30 min WO. Amounts of lysates used for IP were normalized to giantin or NMIIA, as described in Materials and Methods. Red and blue markers indicate BFA and BFA WO samples, respectively. (**B**) Confocal immunofluorescence images of Rab6a colocalization with giantin and NMIIA in LNCaP BFA WO cells. (**C**) Quantification of Pearson’s overlap coefficient for indicated pairs of stained proteins in cells after BFA WO. All confocal images were acquired with same imaging parameters; bars, 10 μm. Data collected from 90 cells of three independent experiments, results are expressed as a mean ± SD; *, p<0.001. (**D**) Giantin and Rab6a W-B of the postnuclear supernatant (PNS) and Golgi membranes collected from the HeLa cells: control and treated with BFA. (**E**) The representative 3D reconstruction of the SIM imaging of Golgi membranes isolated from HeLa cells: non-treated, and treated with BFA followed by incubation with or without Rab6a protein. The regenerated Golgi membranes were stained with giantin; bars, 2 μm. (**F**) Quantification of the average length of Golgi membranes (n=20 from each sample) presented in E. (**G**) Top panel: the reconstructed 3D SIM imaging of Golgi in HeLa cells after 30 min of BFA WO. Cells were stained with Rab6a (green) and giantin (red). The representative area of Golgi is presented in the low panel. Note the Rab6a punctae between giantin-positive emerging Golgi membranes. (**H**) Rab6a, giantin, and NMIIB W-B of complexes pulled down with anti-giantin Ab from the Golgi fraction of LNCaP BFA WO cells pretreated with scramble or SAR1A siRNAs. Non-specific IgG was used as control IP. Lysates containing equal amounts of Rab6a were used for giantin Co-IP.

To validate these screens, we implemented the proof-of-principle experiment, using cell-free reconstitution of Golgi membrane tubule formation (Cluett, de Figueiredo et al., 2016). First, we isolated Golgi membranes from non-treated and BFA-treated HeLa cells according to the protocol established previously(Petrosyan & Cheng, 2013). As shown in **Fig. 3D**, the remnants of the Golgi membranes from BFA-treated samples contain giantin, but not Rab6a. Thus, we use these samples to study the effect of exogenous Rab6a on the fusion of the nascent Golgi membranes. Briefly, the 500 μl of Golgi membranes isolated from BFA-treated HeLa cells were exposed to five μg of active Rab6a protein (Abcam) in presence of the Reaction Mix, which includes 20 μL of 0.5 M EDTA, 10 μL of 100X GTPγS (Cell Biolabs), and 2 μL of ATP (Energy) Regeneration Solution (Enzo). The suspension was incubated in 37 ºC water bath for 15 min. Then, the reaction was stopped by 65 μL of 1 M MgCl_2_. Another sample of Golgi membranes was prepared analogously, except the addition of active Rab6a protein. Next, samples were incubated with anti-giantin Ab followed by Alexa Fluor 488 secondary Ab (1h at RT for each step). Then, samples were gently transferred to the slides, air-dried under the hood in dark, and covered by the glass slips using ProLong antifade mountant (Thermo Scientific). Finally, we reconstructed 3D volume-rendered surfaces from the SIM imaging of Golgi membranes and evaluated the luminal length of these membranes by the ImageJ software. In the samples lacking Rab6a, the largest segregated Golgi structures did not exceed 2 μm, however, the length of Golgi membranes that has been fused in presence of Rab6a was significantly increased and very close to the values we saw in the Golgi fraction from non-treated cells. These data strongly indicate that the maturation of Golgi membranes and their fusion are mediated by the function of Rab6a. Indeed, in another series of 3D SIM imaging of BFA WO HeLa cells, endogenous Rab6a and giantin were found in the close vicinity (**Fig. 3G**, top panel). Remarkably, in many reconstructed Golgi surfaces, we detected Rab6a punctae between the two emerging giantin-stained membranes (**Fig. 3G**, low panel).

Of interest, when BFA WO was performed in cells pre-treated with SAR1A siRNAs, a known blocking connection of Golgi membranes and giantin dimerization(Casey et al., 2016), no association was detected between Rab6a and giantin despite the fact that both have been found in the same Golgi membranes (**Fig. 3H**, top and low panel). However, in cells pre-transfected with scramble siRNAs, the Rab6a dimer was detected with giantin in the same complex. This data imply that the continuous traffic of COPII vesicles is required for the interaction of Rab6a and giantin. Moreover, these results indicate that the interaction between Rab6a and giantin is detectable only in fused Golgi membranes. We also were intrigued that in the nascent Golgi membranes no NMIIA was detected (data not shown), but a reasonable amount of NMIIB was found (**Fig. 3H**, lowest panel), implying that NMIIB interferes with Golgi organization. Importantly, we could not detect NMIIB in the Golgi membranes after SAR1A KD. Altogether, data from **Fig. 3** suggest that during Golgi biogenesis, giantin dimerization and subsequent intercisternal connection are virtually assisted by Rab6a. Meanwhile, we reasoned that PDIA3, the enzyme required for giantin dimerization (Petrosyan et al., 2014), ought to appear at the restored Golgi membranes simultaneously with progression of intercisternal connections. In support of this idea, we found that in LNCaP (c-28) cells recovered from BFA, giantin colocalized with PDIA3. It is important to note that the colocalization points were found mostly on the periphery of adjacent Golgi membranes (**Fig. S3A**, arrows, **and B**), suggesting that this interaction plays a role in connecting Golgi cisternae. In PC-3 and DU145 cells, where Golgi remains fragmented even after BFA WO, the IF signal of PDIA3 at the Golgi membranes was remarkably negligible. Finally, it was also noted that in LNCaP cells the giantin-PDIA3 complex decreased after treatment with BFA, but that this was reversed at 30 min BFA WO (**Fig. S3C**).

These data led us to the next hypothesis, whether a mutual link exists between Rab6a and giantin. In cells treated with scramble siRNAs, after BFA WO the Rab6a IF signal was predominantly compact and perinuclear. The similar distribution of Rab6a was observed in GM130- or GRASP65-depleted cells recovered from BFA; however, in cells treated with giantin siRNAs, after 60 min of BFA WO Rab6a was equally distributed throughout the cell (**Fig. 4A and C**). Quantification of Rab6a IF associated with membranous structures indicated that giantin KD, but not GM130 or GRASP65 KD, blocks the Rab6a recollection into Golgi membranes (**Fig. 4B**). At the same time, cells pretreated with Rab6a siRNAs also failed to recover Golgi, confirming again that Golgi cisternal maturation requires the involvement of Rab6a (**Fig. 4C, D**, right panel, **and E**). **The role for SAR1A and NMIIB in Golgi biogenesis.** The observation of several independent groups strongly indicates that perturbations in COPII vesicle organization lead to structural instability of Golgi. The dominant negative GDP or GTP restricted SAR1 mutants inhibited Golgi recovery from BFA-treated cells (Bannykh, Plutner et al., 2005). Co-knockdown of SAR1(A+B) in HeLa cells induces Golgi scattering; however, it does not impair the ability of Golgi membranes to collect stacks (Cutrona, Beznoussenko et al., 2013). Given that COPII vesicles carry PDIA3 to the Golgi membranes (Petrosyan et al., 2015b), it would be logical to assume that SAR1A depletion has a negative effect on Golgi restoration. Indeed, at 30 min of BFA WO, the vast majority of control cells exhibit either: (a) reformed compact Golgi; or (b) an accumulation of nascent Golgi membranes in the perinuclear zone, depending on the size of cells (**Fig. 4D**, left panel, BFA WO). Conversely, SAR1A-depleted BFA WO cells still demonstrate a lack of perinuclear Golgi (**Fig. 4D**, right panel, **and E**). However, marginal juxtanuclear Golgi was detected in some cells after prolonged (up to 2 hours) recovery (data not shown). These facts prompted us to examine by SIM the giantin distribution in cells after SAR1A depletion. Quantitative analysis of 3D images led us to conclude that cells with SAR1A KD are still able to maintain a limited size Golgi body; however, a great number of giantin-positive punctae were detected in the cytoplasm (**Fig. 4F and G**). Thus, functional SAR1A, i.e., COPII vesicles, appear to be critical not only for giantin dimerization(Petrosyan et al., 2015b) but also for its successful incorporation into the membranous Golgi structures.

**Figure 4.**
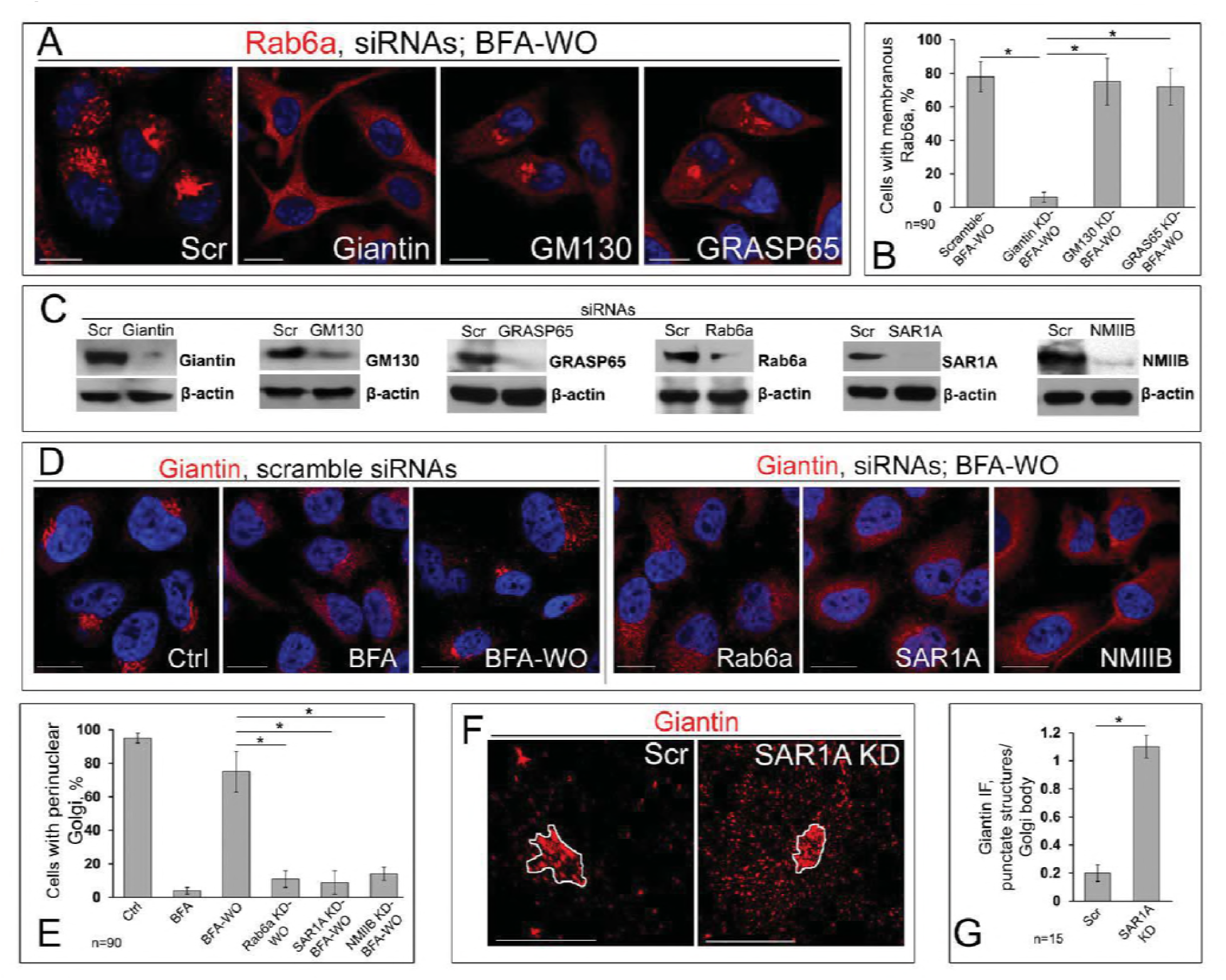
The overlap of giantin and Rab6a during Golgi biogenesis. (**A**) Confocal immunofluorescence images of Rab6a in HeLa cells after BFA WO pretreated with scramble, giantin, GM130 or GRASP65 siRNAs. All confocal images acquired with same imaging parameters; bars, 10 μm. (**B**) Quantification of cells with membranous Rab6a in cells presented in A; n = 90 cells from three independent experiments, results are expressed as a mean ± SD; *, p<0.001. (**C**) Giantin, GM130, GRASP65, Rab6a, SAR1A, and NMIIB W-B of lysates of HeLa cells treated with corresponding siRNAs; β-actin was a loading control. (**D**) Left panel: giantin immunostaining in control, BFA, and BFA WO HeLa cells pretreated with scramble siRNAs. Right panel: giantin immunostaining in BFA WO HeLa cells pretreated with Rab6a, SAR1A, or NMIIB siRNAs. (**E**) Quantifications of cells with perinuclear Golgi in cells from D; n = 90 cells from three independent experiments, results expressed as a mean ± SD; *, p<0.001. (**F**) 3D SIM imaging of giantin in HeLa cells treated with scramble or SAR1A siRNAs. White area indicates the main body of Golgi. (**G**) Quantification of giantin IF integrated intensity (punctate structures/Golgi body) in cells from F; n = 15 cells for each series of 3D imaging experiments, results are expressed as a mean ± SD; *, p<0.001.

It is known that cells pretreated with NMIIA siRNA or its inhibitors demonstrate a significant delay in BFA-induced Golgi disorganization (Duran, Valderrama et al., 2003, Petrosyan et al., 2012b). We have also shown that NMIIA is tethered to Golgi membranes under different drug and stress conditions, such as treatment with EtOH, heat shock, or inhibition of heat shock proteins (HSPs), and depletion of beta-COP (Petrosyan et al., 2016, Petrosyan & Cheng, 2013, Petrosyan & Cheng, 2014). It is also interesting to note that although NMIIA and NMIIB share many biochemical features, they have been found on different membrane domains and ascribed to distinct functions, such as cytokinesis and cell motility (Heimann, Percival et al., 1999, Kelley, Sellers et al., 1996, Maupin, Phillips et al., 1994). In addition, NMIIA and NMIIB remain tightly bound for different lengths of time to F-actin during the ATPase cycle (Kovacs, Wang et al., 2003, Wang, Kovacs et al., 2003). In other words, while the complex of NMIIA and F-actin is the short-term event, the binding of NMIIB to F-actin is prolonged, suggesting NMIIB as a motor protein for generation of sustained tension (Sandquist & Bement, 2010). Importantly, *in vitro* study indicates that NMIIB introduces a twist into the actin filaments, which can cause disintegration of actin bundles into single filaments, and this allows NMIIB to move dynamically back and forth to maintain tension (Norstrom, Smithback et al., 2010). Whether NMIIB interferes with Golgi function, remains enigmatic. It has, however, been proposed that NMIIB is involved in exocytosis and in vesicular trafficking at the *trans*-Golgi and *trans*-Golgi network (Mochida, Kobayashi et al., 1994, Togo & Steinhardt, 2004). In light of these facts, we hypothesize that NMIIA and NMIIB may play a diagonally different role in Golgi morphology. To test this, we performed siRNAs depletion of both NMIIA and NMIIB in HeLa cells. We then measured the Golgi area in mcm^2^, using ImageJ, taking into account only membranous-specific giantin staining. As we predicted, NMIIA KD has no significant impact on Golgi, and Golgi size was comparable to cells transfected with scramble siRNAs (**Fig. S4A-C**). However, in NMIIB-depleted cells, the area of Golgi was significantly enlarged (**Fig. S4A and B**). The data imply that under normal conditions, NMIIB controls Golgi integrity. In support of this view, we then observed that in cells lacking NMIIB, post-BFA recovery of Golgi was significantly impaired (**Fig. 4D**, right panel, **and E**). The detection of NMIIB, but not NMIIA, in the nascent Golgi membranes during their clustering to the perinuclear region (**Fig. 3H**) prompts us to assume that NMIIB, in essence, is a locomotive of Golgi biogenesis. In sum here, the recovery of Golgi requires the action of both COPII vesicles and NMIIB.

### Giantin is essential for post-alcohol Golgi recovery

We have shown recently that EtOH treatment of hepatocytes, both *in vitro* and *in vivo,* results in significant alteration of Golgi morphology (Petrosyan et al., 2015b). This was accompanied by the impaired dimerization of giantin and reduction of its level, but no significant changes in the expression of GRASP65 or GM130 was detected in EtOH-treated cells. Here, for the first time, we analyzed Golgi morphology in liver tissue samples obtained from patients with normal liver function and patients with alcoholic liver cirrhosis. Contrary to the normal cells, most of the cells from alcoholic samples exhibit remarkably altered Golgi structure (**Fig. 5A and B**). Moreover, the level of giantin-dimer in the tissue lysate of these patients was significantly lower than in control samples (**Fig. 5C**). It has been shown that chronic EtOH administration impairs the liver receptor-mediated endocytosis, as in, for example, the uptake of asialoorosomucoid (ASOR) by the asialoglycoprotein receptor (ASGP-R) (Casey, Kragskow et al., 1987). However, internalization and degradation of ASOR were quickly restored upon refeeding by control diet (Sorrell, Casey et al., 1989). This phenomenon prompted us to study post-alcohol Golgi recovery *in vivo*. To do this, we analyzed Golgi morphology in rats feeding with: (a) control diet, (b) EtOH containing (36% of calories) Lieber-DeCarli diet for periods of 5 weeks, and (c) EtOH diet followed by 10 days feeding with the control diet. In control rat hepatocytes, ASGP-R was distributed in Golgi, cytoplasm, and at the periphery of the cell; however, as we have shown before, in hepatocytes from EtOH-fed rats, the cytoplasm was highly vacuolated, and ASGP-R was accumulated in the fragmented Golgi (**Fig. 5D, E, and G**) (Petrosyan et al., 2015b). Of note, in recovered hepatocytes, the number of vacuoles was essentially reduced, Golgi appeared more compact and juxtanuclear, and multiple ASGP-R positive punctae were detected again in the periphery (**Fig. 5F and G**). Golgi recovery was importantly accompanied by giantin re-dimerization and partial restoration of ASGP-R trafficking to the cell surface, as indicated by W-B of both cell lysate and PM fractions isolated from all three categories of rat hepatocytes (**Fig. 5H**).

**Figure 5.**
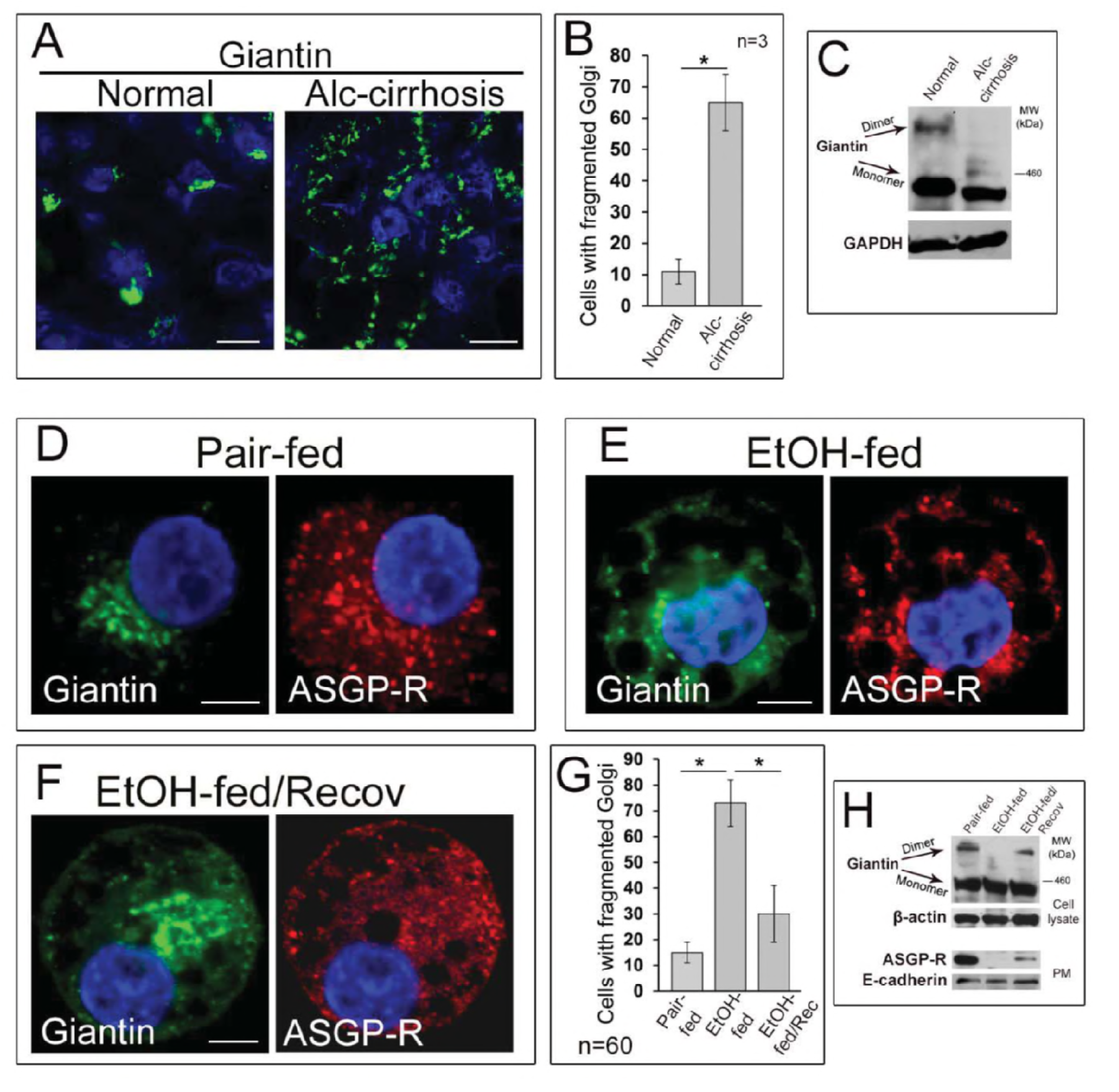
Post-alcohol Golgi biogenesis *in vivo*. (**A**) Confocal immunofluorescence images of giantin in the liver tissue samples obtained from patients with normal liver function and patients with alcoholic liver cirrhosis; bars, 5 μm. (**B**) Quantification of cells with fragmented Golgi; n = 3 samples for each group. Results are expressed as a mean ± SD; *, p<0.001. (**C**) Giantin W-B of lysates from samples described in A. Lysates were normalized to GAPDH. (**D-F**) Giantin and ASGP-R immunostaining in hepatocytes obtained from rats: pair-fed (D), EtOH-fed (E), and EtOH-fed followed by the recovery (F). (**G**) Quantification of cells with fragmented Golgi in cells from D-F; n = 60 cells from two independent experiments, results are expressed as a mean ± SD; *, p<0.001. (**H**) Top panel: giantin W-B of lysates from cells described in D-F; β-actin is a loading control. Low panel: ASGP-R W-B of plasma membrane fractions from cells described in D-F; samples were normalized to E-cadherin.

To examine the precise role of giantin in post-alcohol recovery, we used recombinant HepG2 - VA-13 cells that efficiently express hepatic alcohol dehydrogenase (ADH) (Clemens, Forman et al., 2002). In agreement with our previous observation of these cells (Petrosyan et al., 2015b), we found that treatment with 35 mM EtOH for 72 h induces Golgi fragmentation and reduces the ASGP-R IF signal at the periphery of cells (**Fig. 6A**). By contrast, when cells recovered after EtOH under normal growing conditions for another 48 h, Golgi was restored and ASGP-R signal was consistent with the control sample (**Fig. 6A**, white arrowheads, and **B**). Nevertheless, post-EtOH recovery of ASGP-R peripheral staining was blocked in giantin-depleted cells (**Fig. 6A–C**). Similarly, we found no significant restoration of Golgi and ASGP-R in NMIIB- or Rab6a-depleted cells, nor in cells transfected with dominant negative (GDP-bound) Rab6a(T27N) (**Fig. 6A–C**). Thus, it seems that analogously to BFA WO, post-EtOH Golgi recovery requires giantin, and it is mediated by the GTPase activity of Rab6a and action of NMIIB. As a way to evaluate the trafficking of liver-specific proteins to the cell surface, in addition to ASGP-R, we also measured by W-B the PM content of the polymeric Ig receptor (PIGR) and transferrin. As shown in **Fig. 6D**, the intensity of bands of all three proteins was reduced in EtOH-treated cells, but in EtOH-recovered cells was very close to the value we saw in control cells. Of note, cells recovered from EtOH under giantin, NMIIB or Rab6a depletion, or transfected with Rab6a(T27N), express ASGP-R, PIGR, and transferrin at the level of EtOH-treated cells (**Fig. 6D**). Next, to examine the level of complete glycosylation, we employed the *Sambucus nigra* agglutinin (SNA lectin) that specifically binds to sialic acid attached to terminal galactose in α-2,6 and to a lesser degree, α-2,3 linkage(Cummings & Etzler, 2009). Predictably, in EtOH-treated cells, the PM expression of ASGP-R carrying sialylated N-glycans has been compromised(Casey et al., 2016, Welti & Hulsmeier, 2014), however, the renaissance of Golgi in EtOH-recovered cells was importantly accompanied by the recovery of glycosylation (**Fig. 6E**). Notably, sialylation of ASGP-R was reduced in cells lacking giantin and recovered from EtOH.

**Figure 6.**
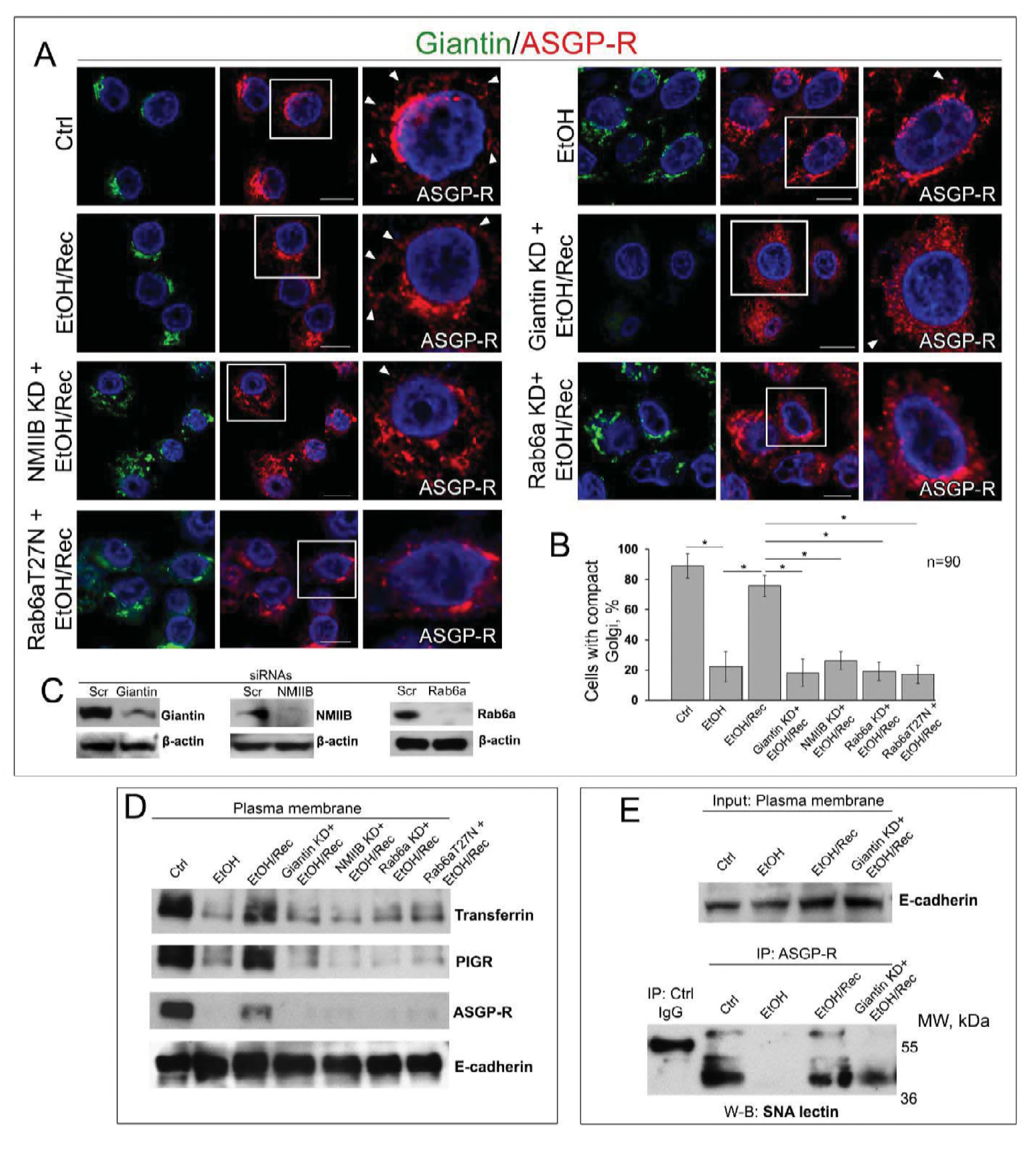
Giantin, Rab6a, and NMIIB are required for *in vitro* post-EtOH recovery of Golgi. (**A**) Confocal immunofluorescence images of giantin and ASGP-R in VA-13 cells: control, EtOH-treated cells, EtOH-treated cells and transfected with scramble, giantin, NMIIB, Rab6a siRNAs, and dominant negative (GDP-bound) Rab6a(T27N) followed by recovery. White boxes are enlarged pictures of ASGP-R presented at the right side. Arrowheads indicate ASPR-R punctae distributed at the periphery of cells. All confocal images acquired with same imaging parameters; bars, 10 μm. (**B**) Quantification of cells with compact Golgi in cells presented in A; n = 90 cells from three independent experiments, results expressed as a mean ± SD; *, p<0.001. (**C**) Giantin, NMIIB, and Rab6a W-B of lysates of VA-13 cells treated with corresponding siRNAs; β-actin was a loading control. (**D**) Transferrin, PIGR, and ASGP-R W-B of plasma membrane fractions isolated from VA-13 cells presented in A; samples were normalized to E-cadherin. (**E**) SNA lectin W-B of ASGPR-IP from the plasma membrane fractions isolated from VA-13 cells: control, EtOH-treated, and recovered from EtOH in absence or presence of giantin siRNAs. The input was normalized to the E-cadherin.

### Is GRASP65 the protein that may compensate the lack of giantin?

Our results so far have suggested that giantin is essential for the biogenesis of Golgi in terms of its compact structure and perinuclear localization. However, in giantin-depleted cells, both GM130- and GRASP65-specific immunostaining do not seem significantly different from the cells transfected with control siRNAs, at least at the level of conventional multifluorescence microscopy (Asante, Maccarthy-Morrogh et al., 2013, Petrosyan et al., 2015b). This suggests that cells that are not under the treatment with Golgi disturbing agent, but are experiencing a deficiency in giantin may launch a compensatory mechanism to maintain Golgi positioning and function. Among Golgi proteins, GRASP65 and GRASP55 are the only two that form stable homodimers to further oligomerize in trans-form (Feng, Yu et al., 2013, Wang, Satoh et al., 2005, Wang, Seemann et al., 2003). It has been proposed that this structural feature of GRASPs is essential to hold adjacent Golgi membranes in stacks: GRASP65 for the *cis*-Golgi, and GRASP55 for the *medial-trans-*cisternae (Pfeffer, 2001, Xiang & Wang, 2010). However, recent observations of GRASP55+65 depleted cells do not completely confirm this concept(Lee, Tiwari et al., 2014); moreover, the siRNA-mediated KD of GRASP65 increases the level of giantin (Petrosyan et al., 2012a). These data are raising the possibility that at the level of *cis-medial-*Golgi, functions of giantin and GM130-GRASP65 may overlap, allowing mutual compensation. In HeLa cells, giantin depletion has no significant effect on GRASP65 dimerization, but the level of GRASP65 tetramer was significantly enhanced (**Fig. 7A and B**). Similar results were obtained from cells treated with NMIIB or Rab6a siRNAs. We reasoned that giantin, and GRASP65 in absence of giantin, may serve as the materials for Golgi intercisternal connections that seem necessary to maintain not only Golgi stacking but also rapid trafficking and processing (Mironov, Sesorova et al., 2017, Rambourg, Clermont et al., 1993, Trucco, Polishchuk et al., 2004).

**Figure 7.**
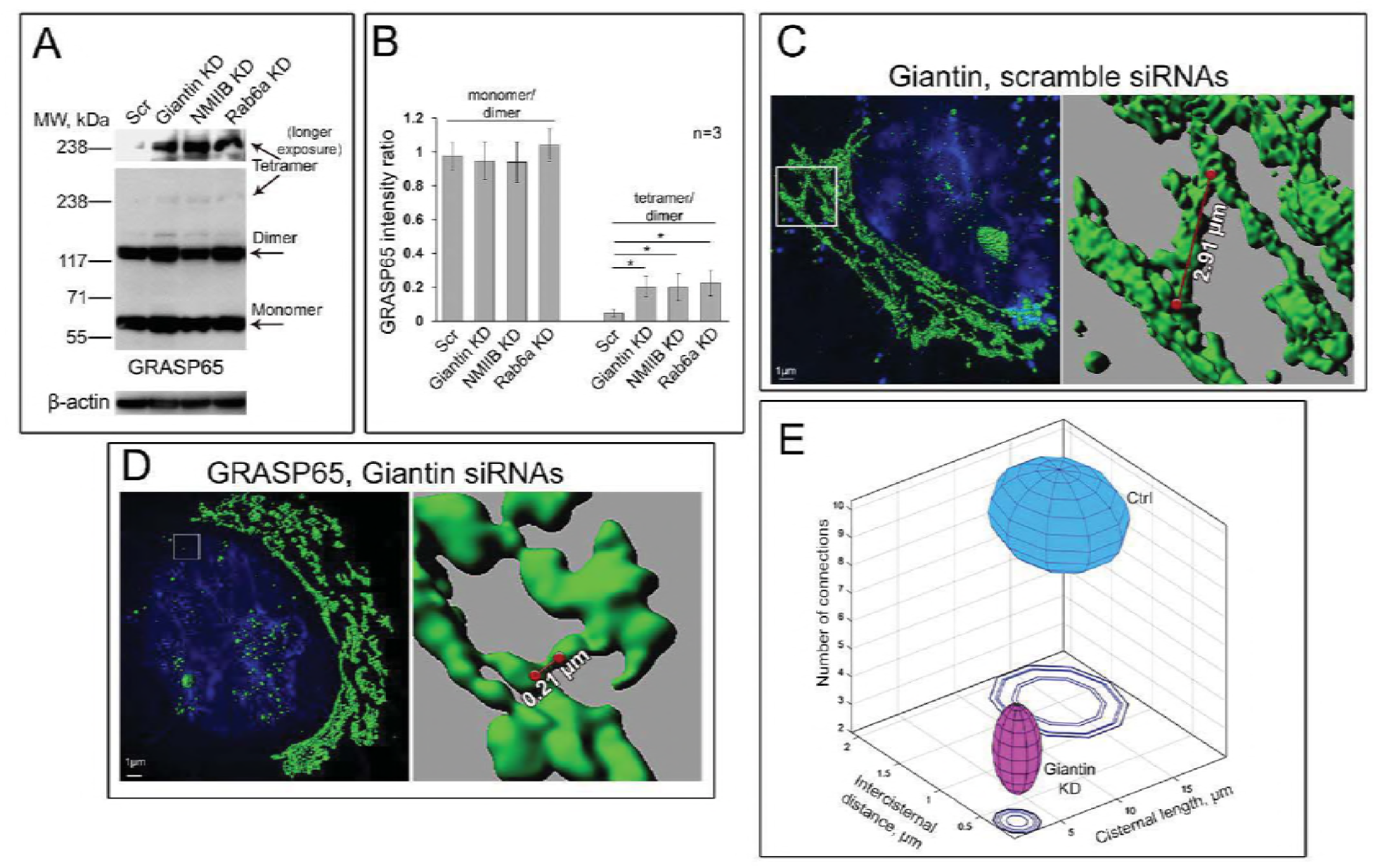
Differential Golgi phenotype in control and giantin-depleted cells. (**A**) GRASP65 W-B of the lysate of HeLa cells: treated with scramble, giantin, NMIIB or Rab6a siRNAs; β-actin was loading control. Longer exposure of GRASP65-tetramer is presented in the top panel. (**B**) Quantification of the intensity of bands corresponding to GRASP65: monomer/dimer and tetramer/dimer. Calculations performed within the same exposure, and data represent mean ± SD from three independent experiments; *, p<0.001. (**C, D**) 3D SIM imaging of giantin and GRASP65 in HeLa cells transfected with a scramble and giantin siRNAs, respectively; bars, 1 μm. Red lines indicate the size of intercisternal connections. (**E**) Statistical analysis of cisternal length, size of intercisternal connections, and their number. Clear separation of observations in control and giantin KD HeLa cells visualized in an XYZ plot where mean and SD were calculated for each parameter (axis) and then used to display an ellipsoid for both conditions. The large ellipsoid represents control data, the small ellipsoid - giantin KD. Contour (shadow) of ellipsoids presented at Z=0. Hypothesis testing for different medians was performed via non-parametric Wilcoxon rank sum test. P-values for length of cisternae, intercisternal distance, and number of connections are 2.2544e-29, 1.2278e-16, 5.2380e-04, respectively with Wilcoxon.

Thus, cells lacking detectable giantin appear to possess Golgi stacks and perinuclear location due to GRASP65 oligomerization. However, several cardinal differences came to our attention after thorough ultrastructural analysis of Golgi performed by a series of SIM imaging. First, average total length of contiguous Golgi cisternae was seen decreased to 1.509 ± 0.609 μm in giantin-KD cells, from 15.12 ± 4.597 μm in control cells. Second, a distance of intercisternal connections was significantly reduced: the “long” communications in control cells (1.509 ± 0.6090 μm) after giantin depletion were replaced by “short bridges” (280 ± 204 nm) (**Fig. 7C and D**). Finally, the number of cisterna-to-cisterna communications per Golgi stack were also lower in cells lacking giantin (4.6 ± 1.578) compared to control cells (8.7 ± 1.636). Kernel density plots calculated for each variable revealed significant differences in the location and shape of the distributions by group (**Fig. S5**). Next, visualization of described differences was combined in an ellipsoid at the XYZ plot (**Fig. 7E**). These results suggest that giantin appears to be critical for maintenance of Golgi cisternae continuity and their connections. **GRASP65 is essential for the Golgi organization in advanced PCa cells**. These findings led us to the conclusion that GRASP65 oligomerization may compensate for the lack of a giantin-dimer. If so, in aggressive PCa cells, where Golgi is depolarized (Petrosyan et al., 2014), GRASP65 oligomerization would then be impaired. To find evidence to support this hypothesis, we first employed 3D SIM imaging to evaluate the level of Golgi disorganization. In addition to PC-3 cells, we used high-passage LNCaP cells (c-123), similar to PC-3 in terms of androgen-unresponsiveness (Iguchi, Fukami et al., 2012, Lin, Hu et al., 2003). In low passage LNCaP (c-28) cells, Golgi is consistent with its classical structure and positioning, and virtually no Golgi fragments were observed. Conversely, in PC-3 cells, and even more in high passage LNCaP (c-123) cells, multiple standalone Golgi elements were detected (**Fig. 8A, Movie S6-8**). Next, in LNCaP (c-123) and PC-3 cells and as predicted, the levels of both giantin-monomer and dimer were reasonably reduced (Petrosyan et al., 2014) (**Fig. 8B**, top panel). As we anticipated, the level of GRASP65 tetramer was reduced in both LNCaP (c-123) and PC-3 cells (**Fig. 8B,** low panel), and instead, the amount of GRASP65-dimer was enhanced, implying that oligomerization of GRASP65, but not its dimerization is impaired (**Fig. 8B and C**). We then asked whether GRASP65-dimer is necessary and sufficient for maintenance of this Golgi phenotype. To gain efficient knockout of GRASP65, we performed its CRISPR/Cas9-mediated deletion in PC-3 cells. In most cells, little if any GRASP65 was detected, confirmed by W-B (**Fig. 8D and E**). These cells demonstrate no indication of Golgi membranous structures, intriguingly suggesting that their structural organization is terminated. However, in cells with detectable GRASP65, Golgi membranes were preserved, as indicated by staining with giantin (**Fig. 8D,** asterisk). Most importantly, we were able to restore fragmented Golgi in GRASP65 KO PC-3 cells, expressing exogenous GRASP65-GFP (**Fig. 8E**). The average size of fragments for rescued Golgi was very close to the values we detected in cells treated with control sgRNAs. Finally, we employed Spearman’s rank correlation coefficient to examine whether dissolution of Golgi is linked to depletion of GRASP65. We found that a positive correlation (R=0.88, p<0.001) exists between the intensity of GRASP65 IF and the size of fragments (**Fig. 8G**). In other words, only cells with residual expression of GRASP65 demonstrated the ability to form membranous structures. Here, we found that in advanced PCa cells, GRASP65 plays a leading role in Golgi organization.

**Figure 8.**
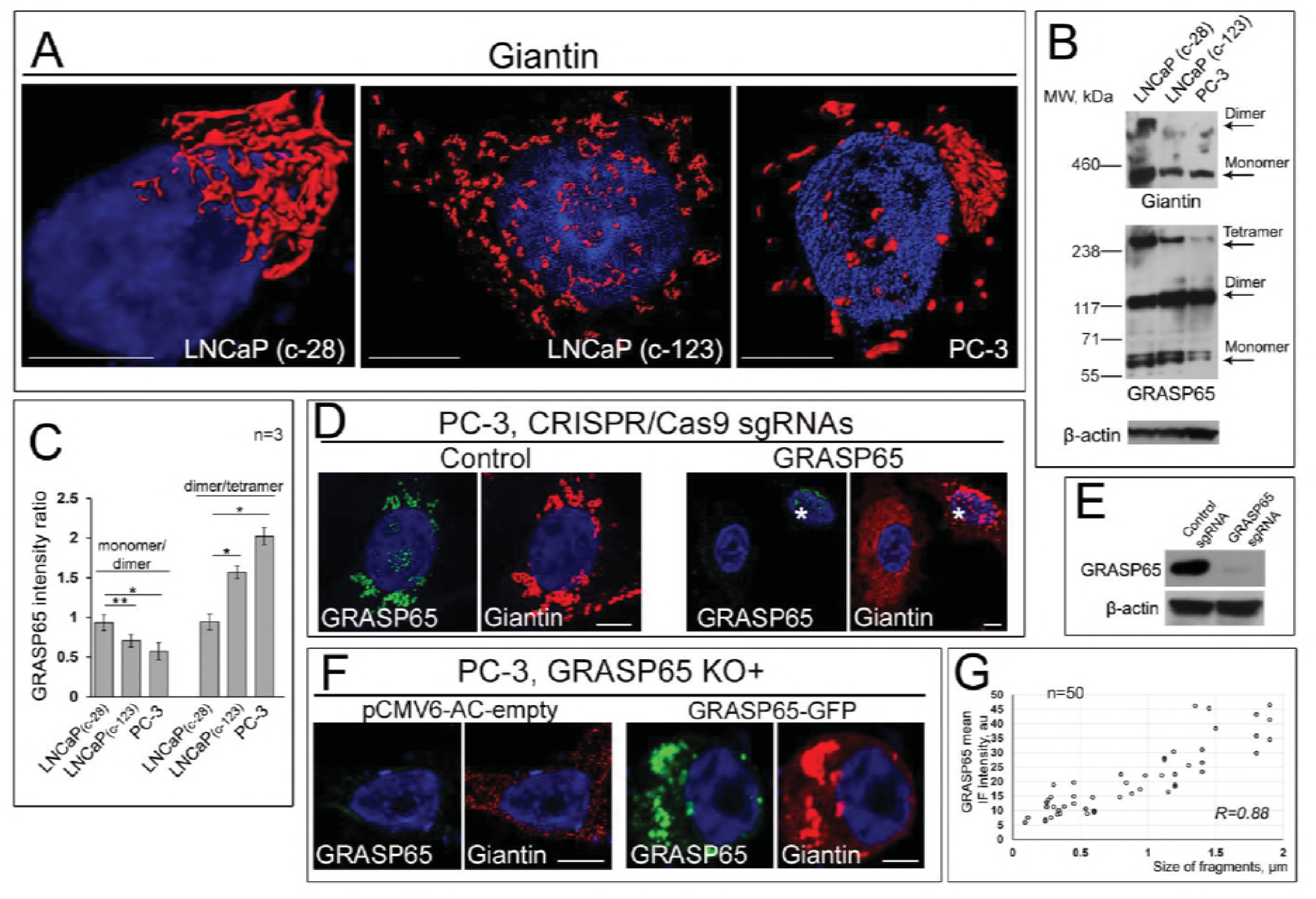
Critical role for GRASP65 in androgen-negative prostate cancer cells. (**A**) 3D SIM imaging of giantin in LNCaP (c-28), LNCaP (c-123), and PC-3 cells; bars, 10 μm. (B) Giantin and GRASP65 W-B of lysates of cells from A; β-actin was a loading control. (C) Quantification of the intensity of bands corresponding to GRASP65: monomer/dimer and dimer/tetramer. Calculations performed within the same exposure, and data represent mean ± SD from three independent experiments; *, p<0.001, **, p<0.01. (**D**) Confocal immunofluorescence images of GRASP65 and giantin in PC-3 cells transfected with CRISPR/Cas9 sgRNAs: control (left panel) and GRASP65 (right panel). All confocal images acquired with same imaging parameters; bars, 10 μm. Asterisk indicates the cell with a residual expression of GRASP65. (**E**) GRASP65 W-B of lysates of cells from D; β-actin was a loading control. (**F**) Confocal immunofluorescence images of PC-3 cells treated with CRISPR/Cas9 GRASP65 sgRNAs followed by transfection with either pCMV6-AC empty vector or GRASP65-GFP; bars, 10 μm. (**G**) Quantification of Spearman correlation coefficient between integrated IF intensity of GRASP65 and diameter of Golgi fragments in fifty PC-3 cells transfected with CRISPR/Cas9 GRASP65 sgRNAs.

## Discussion

The ability of Golgi apparatus to recover after severe attacks is unique and could play a significant role in cellular homeostasis. Here, we describe the role for the largest golgin, giantin, in the maintenance of Golgi stability. Our results clearly indicate that Golgi biogenesis requires giantin dimerization, but that it also depends on the activity of SAR1A, Rab6a, and the action of NMIIB. The appearance of dimerized Rab6a coincides with giantin dimerization and Golgi reconstitution. Importantly, in cells with altered fusion of Golgi membranes Rab6a is not forming complex with giantin. We assume the existence of at least two events that require the virtual involvement of giantin. First, during clustering to the perinuclear region, a giantin and Rab6a monomers from the rims of opposite cisternae are tethering to each other, thus forming dual giantin-Rab6a dimeric complexes (**Fig. 9**). This, in turn, facilitates giantin dimerization, mediated by PDIA3. Intercisternal fusion requires a force that may be created by the action of NMIIB. Although there exists no direct evidence as to how NMIIB tether to the Golgi, our data make this a likely scenario. Second, giantin seems indispensable for the formation of a communication between Golgi stacks. While the function of these internal Golgi “bridges” remains elusive, it was suggested that they could play a role in intra-Golgi trafficking and maintenance of compact shape of Golgi(Mironov et al., 2017, Rambourg et al., 1993, Trucco et al., 2004). The alternative “short” intercisternal communication provided by GRASP65 oligomers seems efficient for Golgi compaction and positioning, but not sufficient for proper processing, because giantin depletion cardinally changes rate and quality of glycosylation (Koreishi et al., 2013). It is important to note here that in normal cells, giantin depletion itself does not lead to the loss of Golgi nucleation because the gradual decline in giantin expression is compensated by the oligomerization of GRASP65. However, giantin appears to be critical for the recovered cells that attempt to restore Golgi architecture after different stresses.

**Figure 9.**
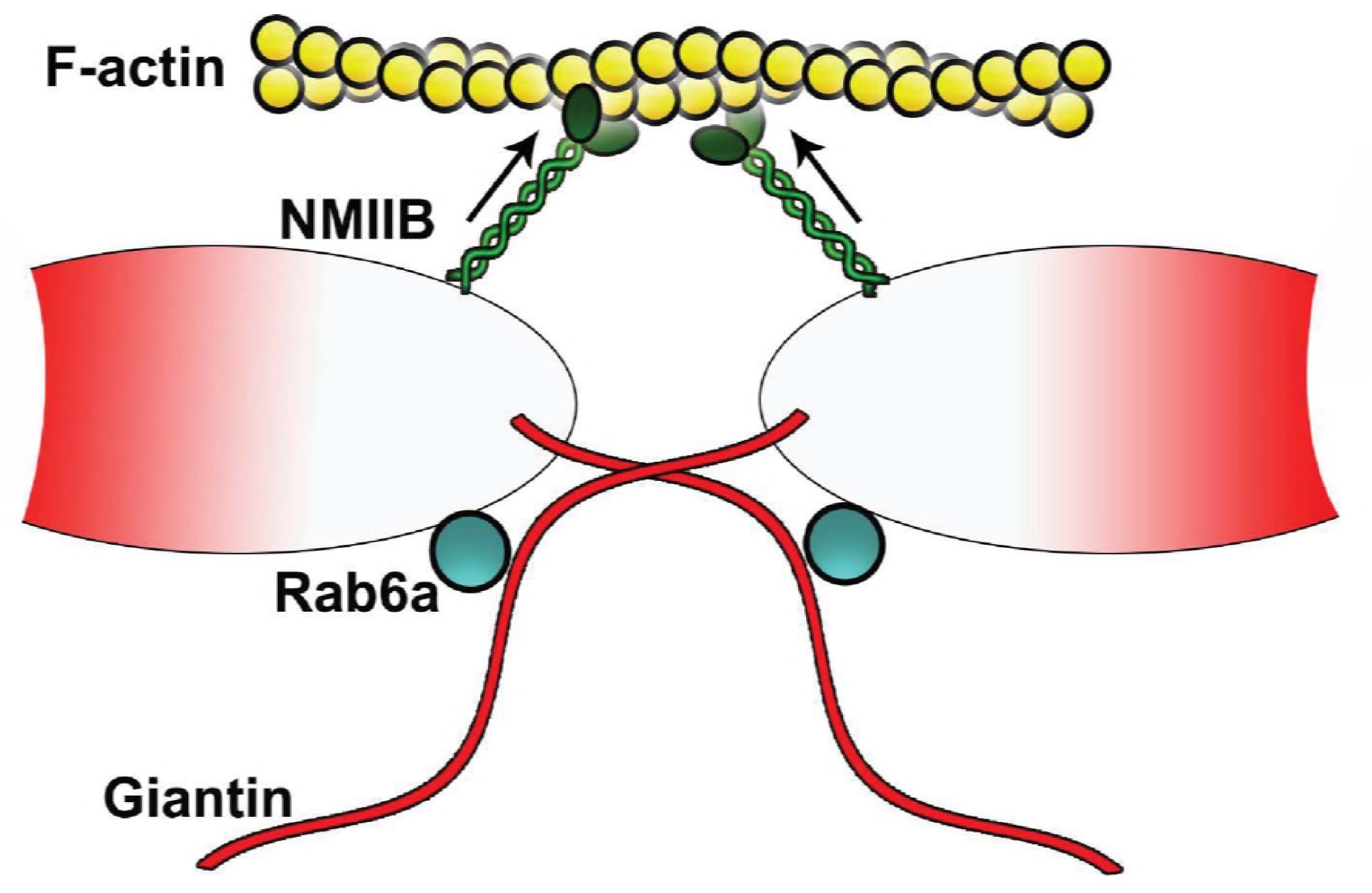
Schematic illustrations of Golgi membrane fusion. Fusion of Golgi membranes is presumably initiated by tethering of giantin from the rim of one cisterna to the rim of opposite cisterna via interaction with Rab6a. This may provide twisting of giantin monomers required for coiled-coil dimeric structure. Then, PDIA3 catalyzes disulfide link in the luminal domain. Such fusion requires the force created by the action of F-actin based motor protein, non-muscle Myosin IIB.

If giantin plays a leading role in the maturation of Golgi membranes, then, what is the role for GM130 and GRASP65, which have been shown to be essential for the lateral cisternal fusion during Golgi assembly and efficient glycosylation(Puthenveedu, Bachert et al., 2006)? Notably, these data were not totally reproduced by another group, which demonstrates that the function of GM130 is rather necessarily for the incorporation of the ER-emanating tubulovesicular carriers into the *cis*-Golgi stacks(Marra, Salvatore et al., 2007). While we do not rule out that under normal circumstances, GM130 and GRASP65 may play a certain role in both events, here we provide evidence that function of the GM130-GRASP65 complex is not essential for the Golgi recovery.

The relationship between the matrix and resident Golgi proteins is increasingly viewed as a mutual alliance, where the function of some may influence the behavior of others (Chang, Hong et al., 2012, Petrosyan et al., 2012a, Puthenveedu et al., 2006, Sengupta et al., 2015, Wei & Seemann, 2010, Xiang, Zhang et al., 2013). On the one hand, golgins and GRASPs are able to form a Golgi matrix framework regardless of the presence of Golgi enzymes (Seemann et al., 2000). On the other hand, Golgi appears as a disassembled structure when some glycosyltransferases fail to localize to the Golgi (Gill, Chia et al., 2010, Pokrovskaya, Willett et al., 2011, Quintero, Valdez-Taubas et al., 2008). For example, a mutation in the membrane-spanning domain of an N-acetylglucosaminyltransferase I caused a dramatic effect on Golgi morphology (Nilsson, Rabouille et al., 1996). In CHO cells lacking N-acetylglucosaminyltransferase V (MGAT5), the Golgi volume density was significantly reduced (Dong, Zuber et al., 2014); however, this was not ascribed to the lack of enzymatic function, given that in the absence (or inhibition) of its precursors, MGAT1 and Mann-II, the Golgi seems unaffected. Still, we do not rule out that this possibility may be limited only to the glycosyltransferases that form complexes with cytoskeletal proteins (Wassler, Foote et al., 2001, Yamaguchi & Fukuda, 1995). It is important to note, however, that the CT of Golgi enzymes plays an essential role not only in their Golgi retention but that it also mediates tethering of NMIIA to the Golgi membranes, thus providing the force for alcohol- or BFA-induced Golgi disorganization (Ali, Chachadi et al., 2012, Petrosyan, Ali et al., 2015a, Petrosyan et al., 2012b, Petrosyan et al., 2016, Petrosyan & Cheng, 2013). Here we provide the evidence indicating that CT of MGAT1 is also required for the Golgi targeting. Since the N-terminal of giantin is projecting into the cytoplasm, it is logical to assume that the direct interaction between MGAT1 and giantin occurs outside of the Golgi. However, we still do not know whether giantin, in addition to the docking function, is also involved in the retention of MGAT1 in the Golgi. This possibility requires further investigations.

Lastly, most golgins and GRASPs have multiple potential N- or O-glycosylation sites, and in case of giantin, this was preliminary confirmed (Petrosyan et al., 2014), suggesting their posttranslational modification may require the assistance of Golgi enzymes. Thus, it is reasonable to conclude that full maturation of Golgi matrix proteins occurs after complete Golgi recovery and the appearance of all resident enzymes at the Golgi membranes. For example, in parasitic protozoan Giardia lamblia, Golgi was not identified in nonencysting cells; nevertheless, the induction of Golgi enzyme activities coincides with stacking of Golgi during differentiation to cysts (Lujan, Marotta et al., 1995). Therefore, our model requires that some of the proteins (which Golgi targeting likely requires giantin) prefer to hold the occupation of membranes until complete recovery of Golgi (**Fig. 1J**). In some aspects, this conception does not fit with the idea that enzymes refill membranes during Golgi biogenesis according to their intra-Golgi (from *trans* to *cis*) location (Jiang et al., 2006, Kasap et al., 2004). However, given that early forming structures in Golgi assembly are inactive in cargo transport (Jiang et al., 2006), the holdup of giantin-sensitive enzymes seem reasonable, because, in this scenario, the accomplishment of Golgi recovery would coincide with the commencement of protein glycosylation.

Our model also suggests that giantin dimerization is a necessity for a successful Golgi biogenesis. Since the formation of giantin-dimer requires functional COPII (Casey et al., 2016) and, broadly speaking, the dynamic of Golgi membranes is linked to the integrity of ER exit site (Marra et al., 2007, Ward et al., 2001), we believe that any substantial disturbance of ER function would inevitably result in Golgi dysfunction and its subsequent disorganization. We show here that in cells lacking giantin, GRASP65 oligomerization may take a leading role in Golgi remodeling and maintenance of its positioning. However, in cells with fragmented Golgi phenotype, GRASP65 consistently fails to form oligomers. Therefore, ER-stress may lead to not only impairment of giantin dimerization, but also to alteration of posttranslational modification of GRASPs and other golgins (Machamer, 2015). In this case, the consequences of Golgi disintegration will appear more severe than one induced by the deficiency in giantin dimerization after treatment with BFA or EtOH. We would thus not exclude the possibility that cancer-specific Golgi fragmentation is influenced by the dysfunction of ER because multiple studies observed the virtual link between ER stress and cancer (Yadav, Chae et al., 2014).

If one assumes that in mammalian cells, compact and perinuclear Golgi is essential for complete glycosylation, does it follow that any disturbance in Golgi morphology will result in the abnormal glycan processing of cargo? The answer lies in the comparison of BFA with other Golgi disruptive agents, such as Cytochalasin D or Nocodazole. Indeed, treatment with Cytochalasin D, an actin destabilization agent (May, Ratan et al., 1998), or Nocodazole, a microtubule depolymerization agent (Turner & Tartakoff, 1989), results in extensive Golgi fragmentation (Petrosyan & Cheng, 2013). Nevertheless, in these cells, the giantin-dependent enzyme, C2GnT-M, still localizes in the Golgi(Petrosyan & Cheng, 2013), suggesting that these chemicals have no significant effect on structure or function of giantin. This could explain the abnormal glycosylation in BFA-treated cells (Irurzun, Perez et al., 1993, Liu, Bastien et al., 2007) that is not found in cells incubated with either Cytochalasin D or Nocodazole (Stallcup, Raine et al., 1983, Xiang et al., 2013). Therefore, while Golgi is anchored in juxtanuclear space *inter alia* by cooperation with cytoskeleton proteins, successful glycosylation is determined by the intra-Golgi location of residential enzymes rather than by the positioning of Golgi (Linstedt, 2004, Rios & Bornens, 2003). The important question of how these separated “mini-Golgi”, induced by either Nocodazole or Cytochalasin D, communicate to maintain glycosylation of cargo remains to be answered. At first glance, clues may come from the vesicular Golgi model rather than a cisternal maturation conception; however, we are still far from a complete understanding of the nature of the many COPI- and COPII-independent vesicular and tubular complexes that have been detected in the Golgi (Cutrona et al., 2013, Matanis, Akhmanova et al., 2002, Petrosyan et al., 2012a, Petrosyan et al., 2012b, Simpson, Nilsson et al., 2006). One of the promising arsenals for these studies is a high-resolution microscopy, without which accomplishment of the current story would have been impossible.

## Materials and Methods

### Antibodies and reagents

The primary antibodies used were: a) rabbit polyclonal – giantin (Novus Biologicals, NBP2-22321), giantin (Abcam, ab24586 and ab93281), ASGP-R1 (Abcam, ab88042), NMIIA (Abcam, ab75590), PDIA3 (Abcam, ab13507), PIGR (Abcam, ab96196), transferrin (Dako, A0061); b) rabbit monoclonal – GAPDH (Cell Signaling Technology, 14C10), Man-I (Abcam, ab140613), MGAT1 (Abcam, ab180578), GM130 (Abcam, ab52649), GRASP65 (Abcam, ab174834); c) mouse monoclonal – NMIIB (Abcam, ab684), GRASP65 (Santa Cruz Biotechnology, sc365434), β-actin (Sigma, A2228), giantin (Abcam, ab37266), SAR1A (Abcam, ab77029), Rab6a (Santa Cruz Biotechnology, sc-81913); d) mouse polyclonal – MGAT1 (Abcam, ab167365), GM130 (Abcam, ab169276). The secondary antibodies (Jackson ImmunoResearch) were: a) HRP-conjugated donkey anti-rabbit (711-035-152) and donkey anti-mouse (715-035-151) for W-B; b) donkey anti-mouse Alexa Fluor 488 (715-545-150) and anti-rabbit Alexa Fluor 594 (711-585-152) for immunofluorescence. Brefeldin A (EMD Chemicals) was dissolved in DMSO immediately before use. Brefeldin A was added to the culture cells at a final concentration of 36 μM, which was followed by incubation at 37°C for the 1 h. To study Golgi recovery after Brefeldin A treatment, cells were rinsed at least 3 times with pre-warmed drug-free medium followed by incubation under regular culture conditions for various durations as described in each experiment. All other chemicals and reagents including EtOH, methanol, and DMSO were of MS-grade/analytical grade and purchased from Sigma.

### Cell culture, EtOH administration, and isolation of rat hepatocytes

HepG2 cells transfected with mouse ADH1 (VA-13 cells) were obtained from Dr. Dahn Clemens at the Department of Veterans Affairs, Nebraska Western Iowa HCS (Clemens et al., 2002). VA-13 cells were grown in Dulbecco’s modified Eagle medium (DMEM) with 4.5g/ml glucose, 10% FBS, non-essential amino acids and 100U/ml of Penicilin+Streptomycin. Twenty-four hours after seeding cells (at ~75% confluence), culture media were changed for one containing 35 mM EtOH for another 72 h. The medium was replaced every 12 h to maintain a constant EtOH concentration. Control cells were seeded at the same time as treated cells and maintained in the same medium; EtOH was replaced by the appropriate volume of medium to maintain similar caloric content. In another series of experiments, after 72 h of treatment with EtOH cells underwent recovery for another 48 h. LNCaP cells are provided by Dr. Ming-Fong Lin at the Department of Department of Biochemistry and Molecular Biology, University of Nebraska Medical Center (Igawa, Lin et al., 2002), and PC-3 and DU145 cells obtained from the American Type Cell Culture (ATCC) grown as described previously.

Primary rat hepatocytes from control and EtOH-fed animals were prepared from male Wistar rats. Rats weighing 140-160 g were purchased from Charles River Laboratories. Initially, animals were fed Purina chow diet and allowed to acclimate to their surroundings for a period of 3 days. Then rats were paired according to weight and fed either control or EtOH containing (35% fat, 18% protein, 11% carbohydrates, 36% EtOH) Lieber-DeCarli diet for 5 weeks (Dyets, Inc). The control diet was identical to the EtOH diet except for the isocaloric substitution of EtOH with carbohydrates (Lieber and DeCarli, 1982). This protocol was approved by the Institutional Animal Care and Use Committee of the Department of Veterans Affairs, Nebraska Western Iowa HCS, and the University of Nebraska Medical Center. For recovery, rats have administered the control diet for 10 days. Hepatocytes were obtained from livers of control and EtOH-fed rats by a modified collagenase perfusion technique as described and used previously by the Casey laboratory (Casey et al., 1987, Tworek, Tuma et al., 1996). The primary hepatocytes isolated from control and EtOH-treated animals were cultured as previously described (Schaffert, Sorrell et al., 2001). Briefly, freshly isolated hepatocytes were seeded in William’s media on collagen coated six well plates with or without coverslips. After 2 hours in culture, cells were washed with PBS, followed by incubation with 5% FBS-Williams media. Cells were maintained at 37°C in 5% CO2 for the indicated time. Additional cell aliquots were washed in cold phosphate-buffered saline, and the pellets were stored at - 70°C for future analysis.

### Human liver tissues

De-identified normal and alcoholic cirrhotic frozen liver tissues were obtained from the Liver Tissue Cell Distribution System (LTCDS), Minneapolis, MN, funded by NIH Contract # HSN276201200017C. Liver tissues were stored at −70 °C until analysis. A portion of liver tissue was homogenized in 0.05M Tris-HCL, 0.25 M Sucrose (pH 7.4) supplemented with protease and phosphatase inhibitors, and centrifuged at 2,200 rpm to obtain post nuclear supernatant (PNS) as previously described (Thomes, Trambly et al., 2012). Proteins from freshly made PNS were subjected to Western blotting for detection of giantin dimerization as described in figure legends.

### Immunoprecipitation (IP) and transfection

For identification of proteins in the complexes pulled down by IP, confluent cells grown in a T75 flask were washed three times with 6 ml PBS each, harvested by trypsinization, and neutralized with soybean trypsin inhibitor at a 2x weight of trypsin. IP steps were performed using Pierce Co-Immunoprecipitation Kit (Thermo Scientific) according to manufacturer instructions. Mouse and rabbit non-specific IgG was used as non-specific controls. All cell lysate samples for IP experiments were normalized by appropriate proteins. To determine whether the target protein was loaded evenly, input samples were preliminarily run on a separate gel with different dilutions of control samples vs. treated, then probed with anti-target protein Abs. The intensity of obtained bands was analyzed by ImageJ software, and samples with identical intensity were subjected to IP. *MYH9* (myosin, heavy polypeptide 9, non-muscle, NMIIA), *MYH10* (myosin, heavy polypeptide 10, non-muscle, NMIIB), *SAR1A*, *GOLGB1* (giantin), *GOLGA2* (GM130), *GORASP1* (GRASP65), *Rab6a*, and scrambled on-targetplus smartpool siRNAs, as well as CRISPR/Cas9 GRASP65 sgRNAs, were purchased from Santa Cruz Biotechnology. All products consisted of pools of three target-specific siRNAs. Cells were transfected with 100 nM siRNAs using Lipofectamine RNAi MAX reagent (Life science technologies). PCMV-intron myc Rab6 T27N was a gift from Terry Hebert (Addgene plasmid # 46782) (Dupre, Robitaille et al., 2006). Transient transfection of cells was carried out using Lipofectamine 3000 (Life Science technologies) following manufacturer protocol. CRISPR/Cas9 Knockout (KO) was performed according to Santa Cruz Biotechnology protocol. PCMV-intron myc Rab6 T27N plasmid was a gift from Terry Hebert (Addgene plasmid # 46782)(Dupre et al., 2006). Rab6a-pCMV3-C-OFPSpark (RFP tag) was ordered from Sino Biological. PCMV6-AC-GRASP65-GFP and GOLGB1 (giantin)-pCMV6-AC-GFP plasmids were obtained from OriGene. Transient transfection of plasmids was carried out using the Lipofectamine 2000 (Life Science technologies) following the manufacturer’s protocol.

### Confocal immunofluorescence microscopy

Staining of cells was performed by methods described previously (Petrosyan & Cheng, 2013). Slides were examined under a Zeiss 510 Meta Confocal Laser Scanning Microscope and LSM 800 Zeiss Airyscan microscope performed at the Advanced Microscopy Core Facility of the University of Nebraska Medical Center. Images were analyzed using ZEN 2009 software. For some figures, image analysis was performed using Adobe Photoshop and ImageJ. Statistical analysis of colocalization was performed by ImageJ, calculating the Pearson correlation coefficient (Dunn, Kamocka et al., 2011).

### Three-dimensional structured illumination (3D-SIM) microscopy and image analysis

SIM imaging of Golgi ribbons was performed on a Zeiss ELYRA PS.1 super resolution scope (Carl Zeiss Microscopy, Germany) using a PCO.Edge 5.5 camera equipped with a Plan-Apochromat 63×1.4 oil objective. Optimal grid sizes for each wavelength were chosen according to manufacturer recommendations. For 3D-SIM, stacks with a step size of 110 nm were acquired sequentially for each fluorophore, and each fluorescent channel was imaged with three pattern rotations with three translational shifts. The final SIM image was created using modules built into the Zen Black software suite accompanying the imaging setup. Analyses were undertaken on 3D-SIM data sets in 3D using IMARIS versions 7.2.2–7.6.0 (Bitplane Scientific). Calculation of intercisternal distances was based on nearest neighbor distances to consider the Nyquist limited resolution, which in our case was around ~94 nm (Kaufmann, Piontek et al., 2012). The 3D mask was obtained by applying a Gaussian filter to merged channels, thresholding to remove low-intensity signals, and converting the obtained stack into a binary file that mapped all voxels of interest for coefficient calculation. For colocalization studies, IMARIS “Colocalization Module” was used. To avoid subjectivity, all thresholds were automatically determined using algorithms based on those of Costes et al. (Costes, Daelemans et al., 2004), based on the exclusion of intensity pairs that exhibit no correlation. Colocalization was determined by Pearson’s coefficient, which represents a correlation of channels inside colocalized regions. After calculation, colocalization pixels were displayed as white. 3D animation was also generated using IMARIS “Animation Module.”

### Isolation of Golgi membrane fractions by sucrose gradient centrifugation

Golgi membrane fractions were isolated using methods described previously (Petrosyan & Cheng, 2013). Cells from ten-to-twelve 75 cm² cell culture flasks were harvested with PBS containing 0.5x protease inhibitors (1.2 ml per flask). Then, after centrifugation for 5 min at 1000 rpm and 4°C, the pellet was resuspended in 3 ml of homogenization buffer (0.25 M sucrose, 3 mM imidazole, 1 mM Tris-HCl; pH7.4, 1 mM EDTA). Cells were homogenized by drawing ~ 30 times through a 25-gauge needle until the ratio between unbroken cells and free nuclei became 20%:80%. The postnuclear supernatant was obtained by centrifugation at 2,500 rpm and 4°C for 3 min and then supernatant adjusted to 1.4 M sucrose by addition of ice-cold 2.3 M sucrose in 10 mM Tris-HCl (pH 7.4). Next, 1.2 ml of 2.3 M sucrose at the bottom of the tube was overlaid with 1.2 ml of the supernatant adjusted to 1.4 M sucrose followed by sequential overlay with 1.2 ml of 1.2 M and 0.5 ml of 0.8 M sucrose (10 mM Tris-HCl, pH 7.4). Gradients were centrifuged for 3 h at 38,000 rpm (4°C) in an SW40 rotor (Beckman Coulter). The turbid band at the 0.8 M/1.2 M sucrose interface containing Golgi membranes was harvested in ~500 μl aliquot by syringe puncture. The fraction at a concentration of ~1.0-1.4 mg protein/ml was used for the experiments mentioned in the Results section.

### Plasma membrane protein isolation and glycan assessment

Plasma membranes were isolated using Pierce Chemical kit (Thermo Scientific) according to their protocol. To analyze glycosylation of proteins from plasma membrane fraction, samples were run on 10% SDS-PAGE followed by W-B with HRP-conjugated *Sambucus nigra* lectin (SNA), which binds preferentially to sialic acid attached to terminal galactose in α-2,6 and to a lesser degree, α-2,3 linkage.

### Statistical analysis

Measurements for the giantin KD do not follow a normal distribution as detected via the Lilliefors test at p <= 0.05 (Cembrowski, Westgard et al., 1979), hence hypothesis testing for different medians was performed via the non-parametric Wilcoxon rank sum test (Gibbons & Chakraborti, 2011) for control versus giantin KD for the three parameters: cisternal length, intercisternal distance, and number of intercisternal connections. Data are expressed as mean ± SD. The rest of the analysis was performed using 2-sided t-test. A value of P < 0.05 was considered statistically significant.

### Miscellaneous

Protein concentrations were determined with the Coomassie Plus Protein Assay (Pierce Chemical Co., Rockford, IL) using BSA as the standard. Densitometric analysis of band intensity was performed using ImageJ.

## Acknowledgments

We thank Dr. Adrian E. Koesters for critical review of the manuscript and Alexander M. Pong for excellent technical assistance. Support for the UNMC Advanced Microscopy Core Facility was provided by the Nebraska Research Initiative, the Fred and Pamela Buffett Cancer Center Support Grant (P30CA036727), and an Institutional Development Award (IDeA) from the NIGMS of the NIH (P30GM106397). This research was supported by the K01AA022979-01 award from the National Institute on Alcohol and Alcohol Abuse (to A.P.), the Nebraska Center for Integrated Biomolecular Communication Systems Biology Core (NIGMS P20-GM113126) (to A.P. and J.R.) and by the Nebraska Research Initiative (to J.R.).

## Author contribution

A.P. designed the study, performed most of the experiments, analyzed the data and wrote the manuscript. S.M. performed some of the experiments, aided in study design. C.C. aided in study design, supervised the ethanol administration, performed isolation of hepatocytes from ethanol-fed rats and aided in manuscript preparation. P.T. aided in the analysis of human samples. J-J. M. and J.C. aided in statistical analysis and manuscript preparation.

